# Suppressing cortical glutamatergic neurons produces paradoxical interictal discharges and seizures

**DOI:** 10.1101/2025.10.02.676593

**Authors:** Shiqiang Wu, Qianwei Zhou, Jaehyeon Ryu, Gen Li, Aditya Iyer, Brian Gill, Hongtao Ma, Hui Fang, Catherine A Schevon, Theodore H Schwartz, Jyun-you Liou

## Abstract

**Introduction:** Seizures are traditionally attributed to excessive excitation or deficient inhibition, yet recent clinical and slice data show they can also paradoxically arise when inhibition outweighs excitation. We chemogenetically suppressed neocortical glutamatergic neurons to test whether such suppression elicits epileptic activity in vivo.

**Methods:** CaMKIIα-driven Gi-coupled hM4Di or Gq-coupled hM3Dq Designer Receptor Exclusively Activated by Designer Drug (DREADDs) were expressed in cortical glutamatergic neurons of Rasgrf2-jGCaMP8m and wildtype C57BL/6J mice. Widefield calcium imaging and 32-channel transparent electrocorticography were performed before and after systemic clozapine-N-oxide (CNO; 1.25–5 mg kg ¹).

**Results:** DREADD activation induced intermittent, large-amplitude, synchronized calcium transients confined to the CaMKIIα-hM4Di focus, accompanied by focal ECoG spikes that persisted for >3 h and sometimes evolved into seizures. By contrast, CNO activation of excitatory neurons via CaMKIIα-hM3Dq DREADDs desynchronized activity without epileptiform discharges. The same process duplicated in wild-type animals provoked similar epileptiform discharges.

**Conclusion:** Selective suppression of excitation can paradoxically drive cortical networks into hypersynchronous, epileptic states, challenging the simple excess-excitation model of ictogenesis. These findings highlight that seizures may result from relative inhibitory dominance, and that considering this ‘excess inhibition’ mechanism could inspire new therapeutic approaches.

## Introduction

Classically, epileptic seizures are understood as resulting from excessive excitation or inadequate inhibition ^1–3^. This imbalance between excitation and inhibition (E/I imbalance) has profoundly shaped our understanding of epilepsy, guiding both research directions and clinical treatment strategies ^4^. For instance, many widely used antiepileptic drugs (AEDs), such as benzodiazepines (*e.g.*, lorazepam, clonazepam) and vigabatrin, exert their therapeutic effects primarily by enhancing GABAergic neurotransmission ^5^, while others such as perampanel ^6^ and topiramate ^7^ act mainly by reducing glutamatergic neurotransmission. The classical E/I model also serves as the theoretical foundation for several emerging therapeutics, ranging from focal transplantation of GABA-releasing interneuron progenitors ^8^ to gene-augmentation strategies that locally boost GABAergic neurotransmission ^9^, as well as network-level neuromodulation modalities that aim to restore E/I balance ^10^.

Although the classical E/I model successfully explains many features of epileptic activity, accumulating experimental and clinical evidence suggests that this conceptualization oversimplifies seizure dynamics ^11,12^. Brain slice studies have revealed that intense inhibitory neuron firing can initiate seizure activity ^13–17^. Clinically, seizures have been reported to emerge paradoxically during states of increased global cortical inhibition, such as non-rapid eye movement (NREM) sleep ^18^, peri-anesthesia periods ^19^, or periods following inhibitory neuromodulation therapies ^20,21^. These striking and paradoxical findings suggest that current therapeutic strategies, broadly aiming to enhance inhibition or diminish excitation, may overlook critical complexities in seizure pathophysiology. This insight holds particular clinical relevance for structural epilepsies, such as those following traumatic brain injury ^22^ and stroke ^23^, where pronounced alterations—loss, dysmorphism or functional disconnection—of cortical glutamatergic circuits are often observed. Recognizing this novel seizure mechanism could inform more nuanced therapeutic approaches, emphasizing personalized treatments tailored to specific seizure pathophysiology and network synchronization profiles rather than uniformly targeting global inhibition or excitation.

In this report, we test whether deliberate focal/regional suppression of neocortical glutamatergic neurons can paradoxically provoke epileptic activity. To do so, we expressed the inhibitory Gi-coupled Designer Receptor Exclusively Activated by Designer Drug (hM4Di-DREADD) selectively in neocortical glutamatergic neurons in several independent mouse lines and silenced them with clozapine-N-oxide (CNO) ^24,25^. Chemogenetic suppression of glutamatergic neurons reliably evoked focal epileptic activities in awake, behaving mice and, occasionally, full-blown focal seizures. To our knowledge, this is the first demonstration that shifting the cortical E/I balance toward inhibition by selectively reducing excitatory drive can itself trigger seizures. Viewed alongside earlier studies in which increased interneuron activity was pro-convulsant ^13–16^, our findings show that epileptic activity can emerge from either side of the E/I ledger, underscoring the need for a framework more nuanced than the classical excess-excitation model.

## Materials and Methods

### Animals

All procedures conformed to National Institutes of Health guidelines and were approved by the Weill Cornell Medicine Institutional Animal Care and Use Committee. Transgenic lines obtained from The Jackson Laboratory were crossed to create the experimental hybrids: Rasgrf2-2A-dCre (JAX #022864), TIGRE2-jGCaMP8m (JAX #037718) and C57BL/6J wild-type (JAX #000664). We employed the hybrid between Rasgrf2-2A-dCre and TIGRE2-jGCaMP8m (hereafter, Rasgrf2-jGCaMP8m) to express jGCaMP8m in superficial cortical glutamatergic neurons. The wild-type mice are used to confirm the reproducibility and to control for potential genetic background effects. Mice of both sexes were employed (20–30g, 8–14 weeks old). Animals were housed individually under a 12 h light/dark cycle with ad libitum food and water; all mice were healthy, immunocompetent and drug-naïve. Animals that did not recover adequately from surgery or were marked as sick by the institutional veterinary staff during routine health checks were excluded from further experimentation.

### Surgery, cranial-window implantation and virus delivery

Animals were selected randomly from the cage at the time of surgery and virus injection; no formal randomisation sequence was generated. All mice were anesthetized with isoflurane delivered in a gas mixture comprising 70% nitrogen (N) and 30% oxygen (O), starting with an induction concentration of 4% and 1%–2% for maintenance. Throughout the procedure, physiological monitoring included heart rate and peripheral oxygen saturation were continuously monitored with a small animal pulse oximeter (Starr Life Science, PA, USA), and their body temperatures were regulated and kept at 37°C using a heating blanket (RWD Life Science, TX, USA). The following parameters were consistently maintained: heart rate ranging from 300 to 450 beats per minute and oxygen saturation exceeding 95%.

Mouse head was secured in a stereotaxic frame and an 8-mm craniotomy that exposes bilateral hemispheres was made, leaving the dura intact. AAV9-CaMKIIα-hM4D(Gi)-mCherry (Addgene viral prep #50477-AAV9, MA, USA; total 5 × 10 vg, 50 nL/min, depth 250–500 µm, lot number: v178325 and v206994) or AAV9-CaMKIIα-hM3D(Gq)-mCherry (Addgene viral prep #50476-AAV9, MA, USA; total 5 × 10 vg, 50 nL/min, depth 250–500 µm, lot number: v175871) was injected at cortical sites (frontal: AP +1.2 mm, ML -1.2 mm; parito-temporal junction: AP -2 mm, ML -1.5 mm) with a microinjector (RWD Life Science, TX, USA). Once the virus injection is accomplished, the cranial window was covered with a sterilized polydimethylsiloxane (PDMS) film (Sylgard® 184, Dow Corning, MI, USA) with or without customized nanomeshed gold-Poly(3,4-ethylenedioxythiophene) polystyrene sulfonate (Au:PEDOT-PSS) electrode array ^26,27^. The PDMS film is sealed with VetBond (3M, MN, USA) and, together with customized stainless steel head plate, was attached to the cranium using C&B Metabond (Parkell, NY, USA). After the surgery, the mice were then housed individually during the 2-week recovery period. Prior to the experiments, the mice underwent a head-fixation adaptation protocol in the imaging chamber for 30 minutes each day over three consecutive days. Experiments commenced at least two weeks post-injection to ensure sufficient DREADD expression and postoperative recovery. All subsequent recordings were performed under awake conditions.

For both AAV9-CaMKIIα-hM4D(Gi)-mCherry and AAV9-CaMKIIα-hM3D(Gq)-mCherry, viral genomes were confirmed by full-length sequencing and aligned to reference plasmids (Addgene #50477 and #50476). No mutations or mis-labeling were detected (see Supplementary Material).

### Activation of DREADD receptors

Clozapine N-oxide (CNO) (SML2304-5MG, MilliporeSigma, MA, USA) was dissolved in 5 ml of 0.9% sterile saline to create a 1 mg/ml solution. Mice received intraperitoneal (i.p.) injections of CNO at a dose ranging from 1.25–5 mg/kg per administration.

### Wide-field optical imaging

Brain fluorescence was imaged with a scientific CMOS camera (Dhyana 400 BSI V3, Tucsen, Fuzhou, China) through a tandem-lens macroscope (Nikon AI-S FX Nikkor 50 mm f/1.2, Tokyo, Japan). Sequential 470-nm and 405-nm excitation was provided by high-power LEDs (M470L4-460/60 and M405L4-405/10, Thorlabs, NJ, USA) and reflected by a dichroic mirror (59022bs, Chroma Technology, VT, USA). Emission was filtered through a 525/50 nm band-pass filter (#86-992, Edmund Optics, NJ, USA). LED illumination and camera exposure were synchronized by an Arduino UNO (Arduino LLC, MA, USA), giving 30 Hz acquisition (24 Hz 470-nm illumination channel, 6 Hz 405-nm illumination channel). Static mCherry images used 565-nm excitation (M565L3-565/70, Thorlabs) and an OSF-512/630 emission filter (FL-004381, Semrock, NY, USA).

### Optical Data Analysis

Optical data were processed in MATLAB R2024b (MathWorks, MA, USA) using custom scripts. First, images were 8-by-8 binned. Subsequently, hemodynamic artifacts were removed by the classic Beer-Lambert law-based method. Widefield frames acquired under 470-nm (calcium-sensitive) and 405-nm (isosbestic) illumination are denoted as *F*_470_ and *F*_405_, respectively ^28^. Baselines, *F*_470,0_ and *F*_405,0_ were estimated with a 3-second sliding window advanced in 1.5-second steps (50% overlap). In each window—and for each pixel independently—the local baseline was defined as the 8th percentile of that pixel’s intensity values within the window. Per-pixel baseline values were linearly interpolated between window centers to obtain continuous baseline traces. The hemodynamically corrected change in calcium fluorescence, Δ*F*/*F*, was computed as equation (1).

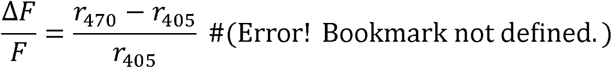

where *r*_470_ = *F*_470_/*F*_470,0_ and *r*_405_ = *F*_405_/*F*_405,0_. To enable comparisons across regions and animals, we further z-scored Δ*F/F* using the mean and standard deviation of Δ*F/F* estimated during the pre-CNO injection period (15 minutes). The resulting single, fixed-referenced neural activity indicator, *Z*, ensures that changes after injection reflect deviations from the same baseline neural activity distribution.

Because there is no consensus, event-level definition of “epileptic discharges” in wide-field calcium imaging, we quantified “epileptiformity” along three complementary axes that reflect canonical features of epileptic population dynamics—semi-periodic bursting, high-gain transients, and hypersynchronous co-activation. The three axes are listed below:

- **Temporal Variance (V)** captures the semi-periodic, fluctuating nature of epileptiform activity. Bursts that recur on sub-second to multi-second scales increase the within-window variance even when waveform shape varies.
- **Peak Amplitude (A)** indexes the intensity of population activation. Epileptiform bursts are marked by large, brief calcium elevations; taking the peak (or high-quantile) within each window yields an amplitude metric that is directly interpretable and sensitive to paroxysmal depolarization–like events.
- **Synchrony Ratio (R)** quantifies spatial hypersynchrony, a hallmark of epileptic population dynamics. R measures the fraction of the imaged ensemble that is co-active within a short temporal window. R thereby distinguishes focal, low-coherence fluctuations from widespread, near-simultaneous recruitment.

All the metrics were defined over a temporal window, *W*, and a local region of interest, *L*. Given a specific time, τ, and a location at the cortical surface, *x*, we defined its temporal window, *W*_τ_, the period starting from τ-2.5 to τ+2.5 seconds and its local region of interest, *L_x_*, as a 600 by 600 microns square region centered at the location, *x*. Subsequently, we define the temporal variance at location *x* and time τ as

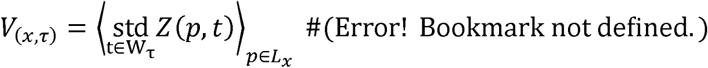

where *p* is the index of pixels. The bra-ket operator stands for spatial averaging.

Analogously, we define the peak amplitude as

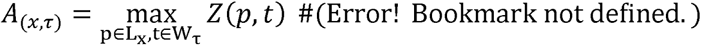

Finally, we define the synchrony ratio, R, as

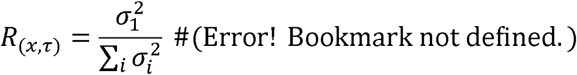

Where σ_*i*_ is the *i*th largest singular value of the matrix *Z*(*p, t*) where *t* ε *W_τ_* and *p* ε *L_x_*. Regions that contain non-brain surface, fibrotic tissues, or significant vasculature were excluded from analysis. To evaluate hemisphere-wide “epileptiformity,” *Z*(*p, t*) was further 10-by-10 binned to speed up computation, and *L_x_* was replaced with the set of all pixels on the hemispheric surface. Subsequent *V, A*, and *R* are estimated analogously.

### Electrophysiology

Electrocorticographic (ECoG) signals were recorded using nanomesh Au:PEDOT:PSS transparent electrode arrays placed under the cranial window. Each array contained 32 electrodes (100 × 100 µm) spaced 1 mm apart within each hemisphere, with the most medial electrode positioned 0.6 mm from the midline. Interconnect wires were 80 µm wide. These arrays permit simultaneous calcium imaging and electrophysiology in the same preparation and were fabricated as described previously ^26^. Signals were amplified with a 32-channel headstage (C3324, Intan, CA, USA) and acquired via an Open Ephys board (SKU: OEPS-9029, Lisbon, Portugal) at 2000–4000 Hz, with a 500 Hz anti-aliasing filter applied at the amplifier level.

For wild-type mice, ECoG signals were recorded via a normal saline-filled glass electrode, subsequently amplified 1000× and band-pass filtered between 1 and 500 Hz (AM Systems Model 1800, WA, USA). The filtered signals were digitized using a CED Power 1401 interface and acquired on a PC running Spike2 software (Cambridge Electronic Design, Cambridge, UK).

### Electrophysiology Data Analysis

Electrode channels with impedance >1 MΩ at 1 kHz were excluded from analysis. ECoG signal preprocessing and spike detection were performed by first subtracting the average across all channels from each channel, followed by passing the signals through a zero-phase shift, 4^th^-order Butterworth bandpass filter. The Hilbert transform was then applied to extract the signal envelope. Interictal epileptiform discharges (IEDs) were detected from the envelop using MATLAB’s build-in findpeaks function, with parameters including a minimum peak width of 20 ms, a maximum peak width of 70 ms ^29^, a minimum peak height of 300 μV,a maximum peak height of 1,500 μV (to exclude artifact), a minimum peak prominence of 200 μV, a minimum peak distance of 100 ms ^30^. All detected peaks were further reviewed by two physicians, JYL and SQW, to exclude artifacts. Line length, a standard measure of ECoG activity, was calculated as the sum of absolute differences between consecutive samples within a 1-s sliding window.

### Data availability

The data sets are available from the corresponding author on reasonable request.

## Results

### Calcium imaging reveals focal network instability after chemogenetic suppression of cortical excitatory neurons

Figure 1A summarizes the experimental workflow. Each mouse was prepared with an 8-mm cranial window sealed by PDMS and overlaid with a nanomesh Au:PEDOT-PSS ECoG array (Figure 1B), permitting simultaneous wide-field calcium imaging and electrophysiology. A focal injection of AAV9-CaMKIIα-hM4Di-mCherry into the parietal-temporal junction of the right hemisphere enabled chemogenetic suppression of local cortical CaMKIIα^+^ glutamatergic neurons ^31,32^.

**Figure 1:**
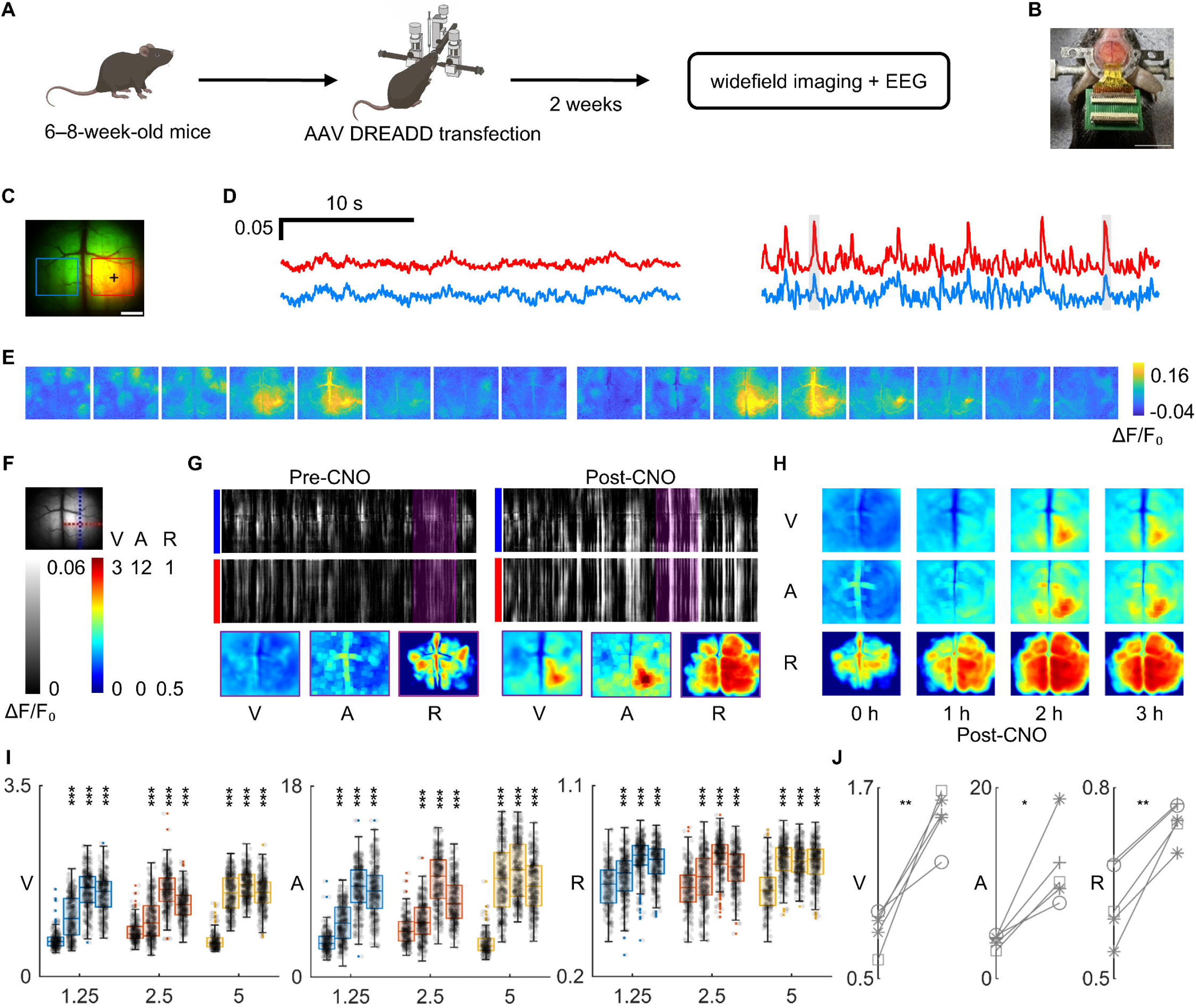
Chemogenetic Modulation of Cortical Activity. (A) Schematic of the experimental workflow. Mice underwent stereotaxic injections of DREADD-carrying AAVs, followed by an 8-mm craniotomy, PDMS window implantation, and EEG/ECoG electrode placement. (B) Post-surgical setup showing the position of the ECoG electrodes on the cortical surface. Scale bar= 1 cm. (C) Representative fluorescence images show successful AAV transfection, evidenced by mCherry expression (red) at the injection site and GCaMP (green) across the cortex. ‘+’ indicates the virus injection site. Two regions of interest (ROIs) were selected: the virus-injected region (Inj. Region, red) and a contralateral control region (Ctrl. Region, blue). Scale bar = 2mm. (D) Representative wide-field optical imaging (WFOI) traces showing normalized ΔF/F signals from the virus-injected region (Inj. Region, red) and contralateral control region (Ctrl. Region, blue) at baseline and after CNO administration. Colors correspond to ROIs defined in the fluorescence image. (E) Heatmaps of ΔF/F activity correspond to the two gray shaded periods in (D), revealing hyperactivity in the Inj. Region. (F) Schematic of analysis lines overlaid on the dorsal cortical surface. The red and blue dashed lines each represent a 1-mm-wide region used for extracting spatiotemporal ΔF/F signals in orthogonal orientations—blue (sagittal) and red (coronal), respectively. The intersection of these lines marks the site of the virus injection. (G) Representative spatiotemporal ΔF/F activity before (left) and after (right) CNO injection (5 mg/kg, i.p.). Top row: ΔF/F line-scan images (30 s) extracted along the vertical blue line in F (sagittal orientation). Middle row: ΔF/F line-scans along the horizontal red line in F (coronal orientation). Bottom row: 5-second averaged heatmaps (purple box in upper panels) showing temporal variance (V), peak amplitude (A), and synchrony ratio (R), respectively. (H) Group-averaged maps of V, A, and R at baseline (0 h) and during the first, second, and third hour following CNO injection (5 mg/kg, i.p.). Each map reflects the first 15 minutes of the respective hour. (I) Quantification of V, A, and R during the first 15 minutes of each hour within 3 hours post-CNO (1.25, 2.5, and 5 mg/kg), compared to baseline, which is also 15-minute long. Each dot represents a 5-second time window. Boxes show the median and interquartile range (IQR), and whiskers extend to the most extreme data points not considered outliers. \*\*\**P* < 0.001, two-tailed Mann–Whitney U test compared to the pre-CNO period. (J) Paired comparisons of V, A, and R between baseline and the second hour after CNO injection (first 15 minutes, 5 mg/kg, n = 5 mice). Each line connects data from the same mouse. \*\**P* < 0.01 for V and R; \**P* < 0.05 for A, paired t-test.

We first examined Rasgrf2-jGCaMP8m mice. This transgenic line was chosen because only a sparse subset of layer 2/3 cortical neurons express jGCaMP8m, thereby minimizing the risk of GCaMP-related network hyperexcitability previously reported in lines with widespread cortical expression ^33–35^. Robust mCherry fluorescence at the injection core verified successful viral transfection (Figure 1C).

Following a single intraperitoneal dose of CNO (5 mg/kg), large-amplitude calcium transients emerged in the injected hemisphere (Figure 1D-E). To characterize these changes spatially, we defined two orthogonal, 1-mm-wide regions of interest centered on the injection site (blue dashed line, sagittal orientation; red dashed line, coronal orientation; Figure 1F). Spatiotemporal ΔF/F_0_ signals extracted along these lines (Figure 1G) revealed marked alterations in activity after CNO administration.

Because there is no universally accepted calcium-imaging criterion for identifying epileptiform events, we quantified calcium fluorescence signals using three complementary metrics that capture hallmark features of epileptic neural activity: temporal variance (*V*), which reflects the semi-periodic alternation between quiescence and bursts of intense firing; peak amplitude (*A*), which indexes the strength of population activity; and synchrony ratio (*R*), which measures the degree of co-activation across cortical locations. All three measures increased prominently in the virus-injected region after CNO administration (Figure 1G, bottom panels).

Group-averaged maps of *V*, *A*, and *R* at baseline and over the three hours following CNO (5 mg/kg) revealed sustained and spatially localized increases in all three parameters (Figure 1H). Across three CNO doses (1.25, 2.5, and 5 mg/kg), quantitative analysis confirmed significant elevations in V, A, and R relative to baseline (two-tailed Mann–Whitney U test, *P* < 0.0001 for all; Figure 1I). Paired comparisons between baseline and two hours after CNO injection (5 mg/kg; *n* = 5 mice) showed significant increases in V (*t*(4) = –5.78, *P* = 0.0045), A (*t*(4) = –4.14, *P* = 0.0143), and R (*t*(4) = –5.80, *P* = 0.0044) (Figure 1J). Together, these data demonstrate that chemogenetic suppression of cortical CaMKIIα^+^ glutamatergic neurons reliably induces focal epileptic neural activity detectable with temporospatial calcium-imaging metrics.

### Chemogenetically activating glutamatergic neurons does not produce epileptic neural activity

To exclude the possibility that increases in V, A, and R after CNO were attributable to nonspecific drug effects or viral expression—and to probe the directionality of the manipulation—we repeated the imaging protocol using AAV2/9-CaMKIIα-hM3Dq-mCherry (chemogenetic activation of glutamatergic neurons) under identical conditions in a separate mouse. In contrast to hM4Di, CNO (1.25–5 mg/kg) did not increase any metric within the injected cortex. Instead, we observed reductions in *V* and *A*, while changes in *R* were inconsistent (Figure 2). Spatiotemporal Δ*F*/*F*LJ line scans and group-averaged maps corroborated the absence of aberrant activity. These controls indicate that CNO and AAV expression alone are insufficient to account for the effects in Figure 1 and, critically, that the paradoxical hyperexcitability is specific to suppressing (hM4Di), not activating (hM3Dq), CaMKIIα^+^ glutamatergic neurons, and not due to CNO off-target effects.

**Figure 2:**
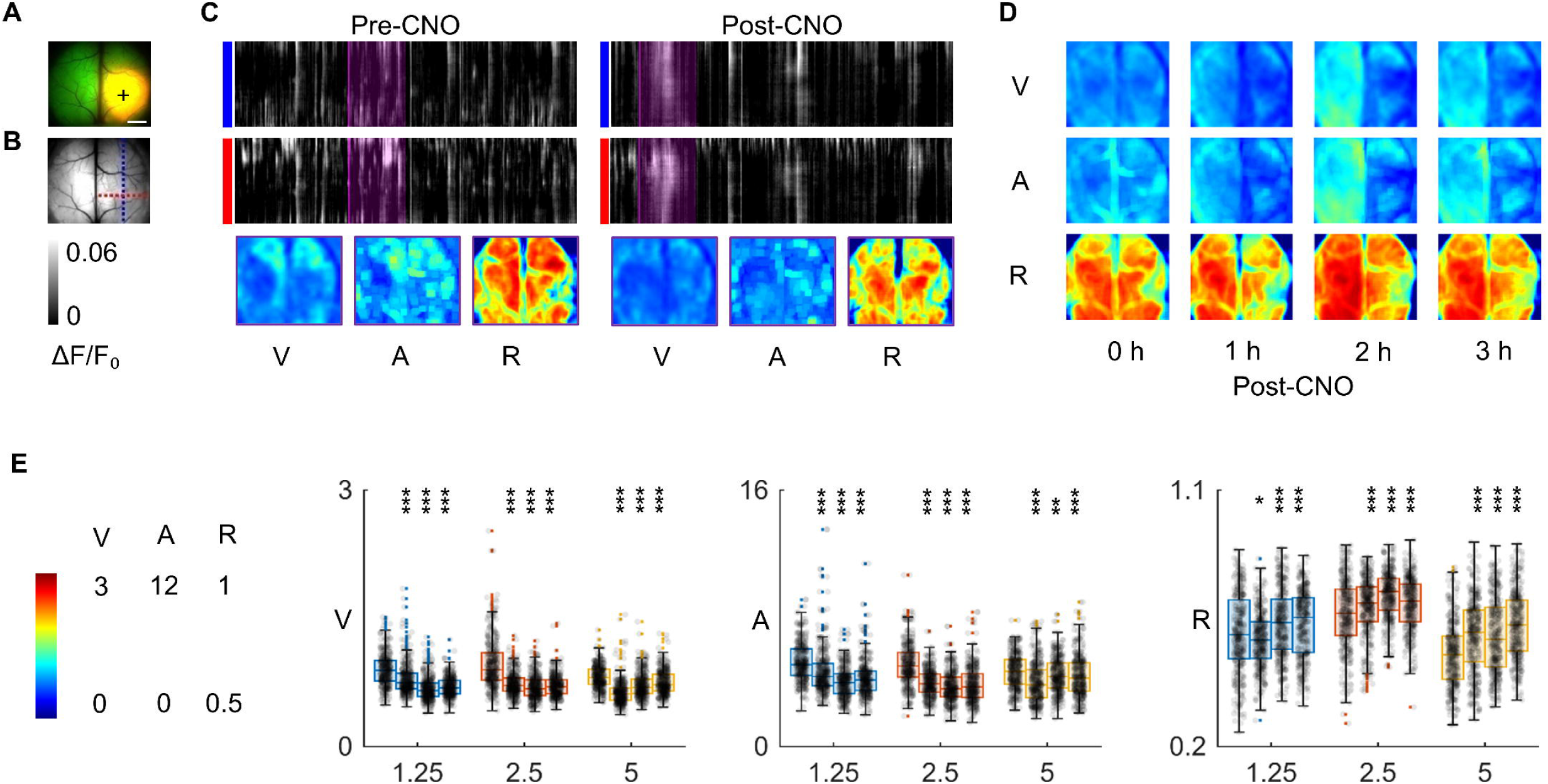
Control experiments using AAV2/9-CaMKIIα-hM3Dq-mCherry. (A) Representative fluorescence image showing mCherry expression (red) in the virus-injected region and GCaMP fluorescence (green) across the cortex. Scale bar = 2 mm. (B) Schematic illustration of the analysis grid overlaid on the cortical surface. The red and blue dashed lines represent 1-mm-wide ROIs for extracting horizontal and vertical line-scan ΔF/F signals, respectively. (C) Spatiotemporal ΔF/F traces extracted from the vertical (top) and horizontal (middle) ROIs before (left) and after (right) CNO injection (5 mg/kg, i.p.). Lower panels show 5-second averaged ΔF/F heatmaps from the highlighted epochs (purple boxes) used for quantifying temporal variance (V), amplitude (A), and synchrony ratio (R). Unlike hM4Di-expressing mice, no prominent post-CNO increases in any metric were observed. (D) Group-averaged V, A, and R maps at baseline (0 h), and at 1 h, 2h, and 3h post-CNO injection. (E) Quantification of V, A, and R during the first 15 minutes of each hour within 3 hours post-CNO (1.25, 2.5, and 5mg/kg), compared to baseline. Significant reductions were observed in V and A at all doses, while changes in R were variable across conditions. (*P < 0.05, **P < 0.01, ***P < 0.001, two-tailed Mann–Whitney U test).

### Electrocorticography confirms that calcium-imaging changes correspond to interictal epileptiform discharges

Because calcium fluorescence signals, while often used as proxies for network events, are not themselves the gold standard for defining epileptiform discharges, we performed simultaneous ECoG recordings with transparent electrode arrays (Figure 3). This allowed us to verify that the large-amplitude calcium transients observed after CNO corresponded to bona fide interictal epileptiform discharges.

**Figure 3:**
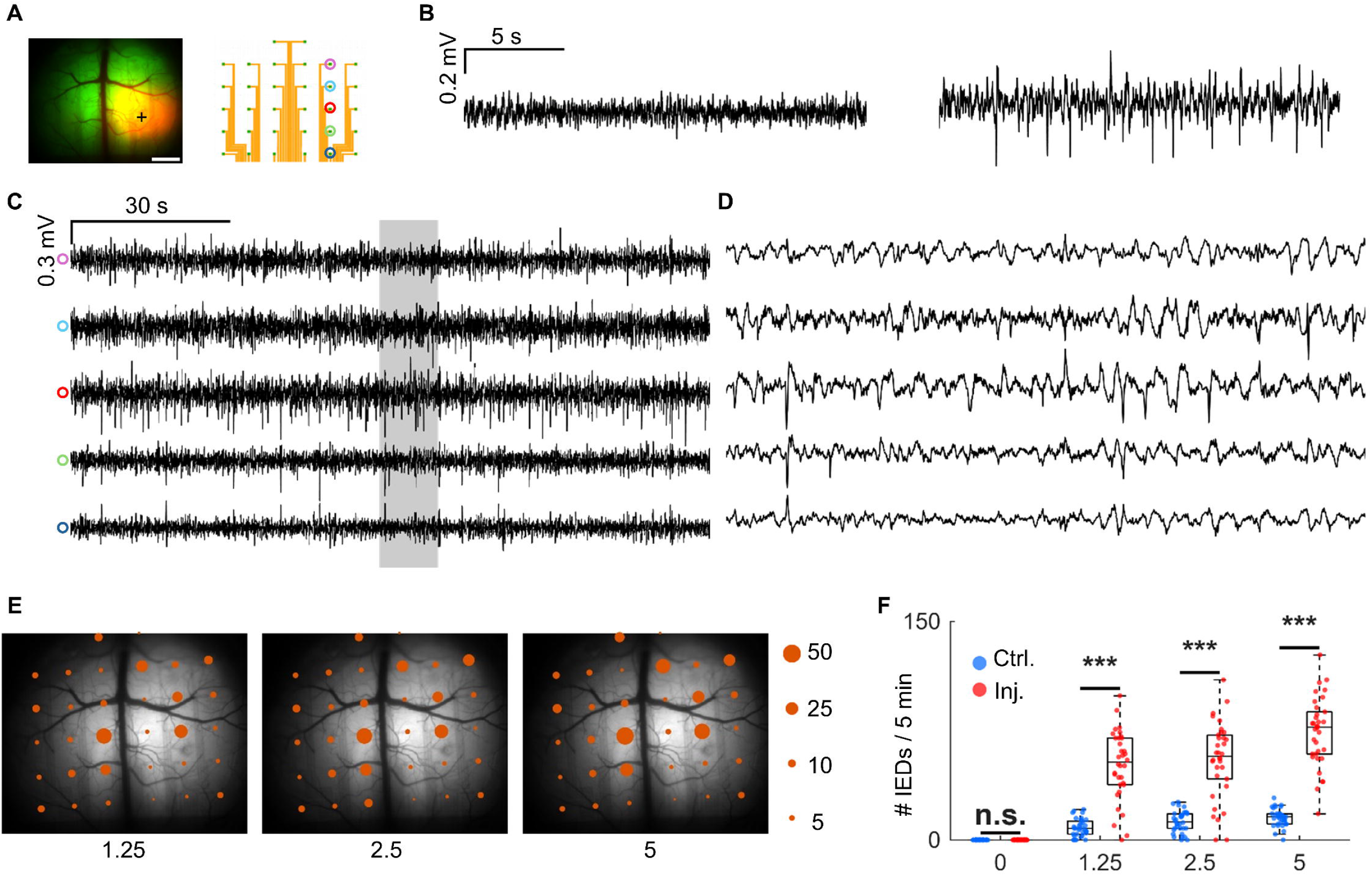
EEG Analysis Reveals Localized Epileptiform Activity Following Chemogenetic Manipulation. (A) Spatial layout of the 8-mm ECoG electrode array over the cortex. Scale bar = 2mm. (B) Representative EEG traces from electrodes surrounding the injection area before (left) versus after (right) CNO injection. (C) Representative EEG traces from five channels within the same electrode column, each preceded by a uniquely colored circle indicating the channel identity. The red circle denotes the channel at the injection site. Channels near the injection site show more frequent spiking activity compared to more distant sites. (D) A zoomed-in view of the gray-highlighted window in (C), illustrating detailed spike morphology across the five channels. (E) Spatial mapping of epileptic spike activity across the cortex following IP injection of CNO at three concentrations (1.25, 2.5, and 5 mg/kg). Each orange circle represents the average number of epileptiform spikes per 5 minutes recorded from the corresponding ECoG electrode site. Circle size is proportional to spike count, as indicated in the legend on the right. (F) Dose-response relationship between CNO concentration and IED activity. Spike counts per 5-minute recording from injected (red) and contralateral (blue) cortices under increasing CNO doses (0, 1.25, 2.5, 5 mg/kg, i.p.). Asterisks indicate significant differences between hemispheres at the same dose (two-sample t-test). Within each hemisphere, spike counts significantly increased at all doses compared to baseline (*P* < 0.0001). Dose–dose comparisons showed significantly higher spike counts at 5 mg/kg versus 1.25 and 2.5 mg/kg in the injected hemisphere (*P* = 0.0002 and 0.0024, respectively). Smaller dose–dose differences were observed in the contralateral cortex. A significant positive dose–response correlation was confirmed by Spearman’s rank test in both hemispheres (ρ = 0.368–0.404, 95% confidence interval, *P* < 0.001).

In Rasgrf2-jGCaMP8m mice expressing hM4Di, CNO administration (5 mg/kg) induced repetitive IEDs — brief, stereotyped spikes or sharp waves — localized to electrodes near the viral injection site, whereas baseline traces were normal (Figure 3B). A column of five electrodes spanning the injection core revealed a spatial gradient in discharge amplitude, peaking centrally and diminishing with distance (Figure 3C–D).

Spatial maps of IED frequency showed a focal distribution centered on, but not strictly limited to, the injection site, with activity remaining spatially restricted and increasing with CNO dose (1.25, 2.5, and 5 mg/kg; Figure 3E). Quantitatively, the injected hemisphere also showed significantly higher IED counts than the contralateral hemisphere at each dose (two-sample t-test, *t*(66) = -10.331, *P* < 0.0001, *t*(66) = -9.111, *P* < 0.0001, and *t*(66) = -14.059, *p* < 0.0001, respectively). All CNO doses produced significantly higher IED counts compared to baseline in the injected hemisphere (*t*(66) = -12.769, -12.225, and -18.330, respectively; all *p* < 0.0001, two-sample t-tests). Within the injected hemisphere, IED counts increased monotonically with CNO concentration, with a significant positive dose–response correlation (Spearman’s ρ = 0.368, P < 0.001; Figure 3F).

These findings confirm that the network changes detected optically in Figure 1 correspond to electrophysiologically defined interictal epileptiform discharges, and that their frequency scales with the level of chemogenetic suppression of cortical CaMKIIα^+^ glutamatergic neurons.

### Chemogenetic-induced hyperexcitability is conserved across cortical regions and genetic backgrounds

To test whether the effect observed in parietal cortex (Figure 1) was region-specific, we instead targeted the frontal cortex in Rasgrf2-jGCaMP8m mice (Figure 4A). CNO administration again produced spatially restricted increases in calcium activity confined to the injection site (Figure 4A–C). Group-averaged maps of *V*, *A*, and *R* showed consistent post-CNO elevations in the targeted cortex regardless of injection location (Figure 4D). Quantitative analysis across three CNO concentrations (1.25, 2.5, and 5 mg/kg) revealed a significant overall trend toward increased *V*, *A*, and *R* relative to baseline in the injected hemisphere (two-tailed Mann–Whitney U tests, except *R*-5 mg/kg, *P* < 0.0001 for all conditions; Figure 4E).

**Figure 4:**
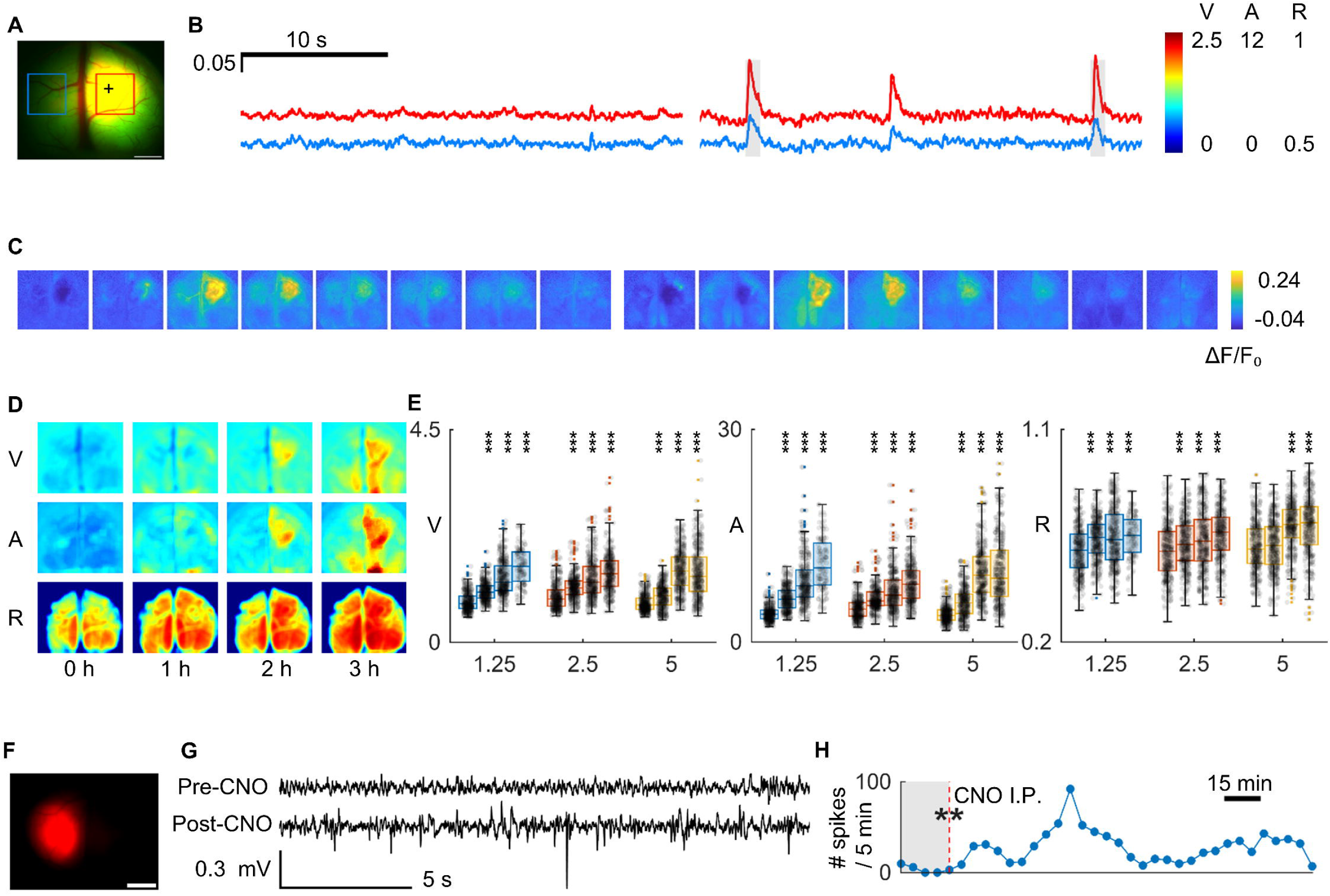
Seizure Activity Across Different locations and Mouse Models. (A) Representative fluorescence images show mCherry expression (red) at the injection site and GCaMP (green) across the cortex. Two regions of interest (ROIs) were selected: the virus-injected region (Inj. Region, red) and a contralateral control region (Ctrl. Region, blue). Scale bar = 2mm. (B) Representative wide-field imaging traces showing normalized ΔF/F signals from the virus-injected region (Inj. Region, red) and contralateral control region (Ctrl. Region, blue) at baseline and after CNO administration. Colors correspond to ROIs defined in the fluorescence image. (C) Heatmaps of ΔF/F activity correspond to the gray shaded area in (B), revealing hyperactivity in the Inj. Region. (D) Averaged heatmaps of V, A and R at baseline (0 h) and during the first, second and third hour after CNO injection (first 15 min of each hour). (E) Quantification of V, A, and R during the first 15 minutes of each hour within 3 hours post-CNO (1.25, 2.5, and 5 mg/kg), compared to baseline, as shown in Figure 1I.\*\*\**P* < 0.001; n.s., *P* > 0.05; two-tailed Mann–Whitney U test compared to the pre-CNO period. (F) Viral expression pattern in wild-type mice. Red regions highlight areas of virus-mediated expression. Scale bar=2mm. (G) Representative EEG traces from wild-type mice before (top) and after (bottom) CNO injection. At baseline, recordings appear normal; however, following CNO administration, distinct epileptiform spike activity emerges. (H) Time course of spike frequency after CNO injection in a wild-type mouse (5-min bins). Spike counts significantly increased after drug administration compared to baseline (*P*=0.0056, Wilcoxon signed-rank test).

We next asked whether chemogenetic suppression alone—without any genetically encoded calcium indicator—could produce detectable electrophysiological correlates. Again, this was motivated by prior reports that some calcium indicator lines can themselves increase cortical excitability (Steinmetz et al., 2017). In wild-type C57BL/6J mice expressing hM4Di-mCherry, robust mCherry fluorescence confirmed viral expression in the targeted cortex (Figure 4F). Baseline ECoG recordings were normal, but after CNO injection, repetitive IEDs emerged and persisted for several hours (Figure 3G). Quantification showed a significant increase in IED frequency compared to baseline (Wilcoxon signed-rank test, *P* = 0.0056; Figure 4H).

### Chemogenetic suppression can trigger focal-onset seizures with secondary generalization

Beyond interictal activity, chemogenetic suppression of cortical CaMKIIα^+^ glutamatergic neurons occasionally escalated to overt seizures. Across all recording sessions we detected three seizures in two mice; one of which is shown in Supplementary Video 1 and Figure 5, illustrating a representative episode spanning pre-ictal changes, ictal onset, propagation, termination, and post-ictal suppression.

**Figure 5:**
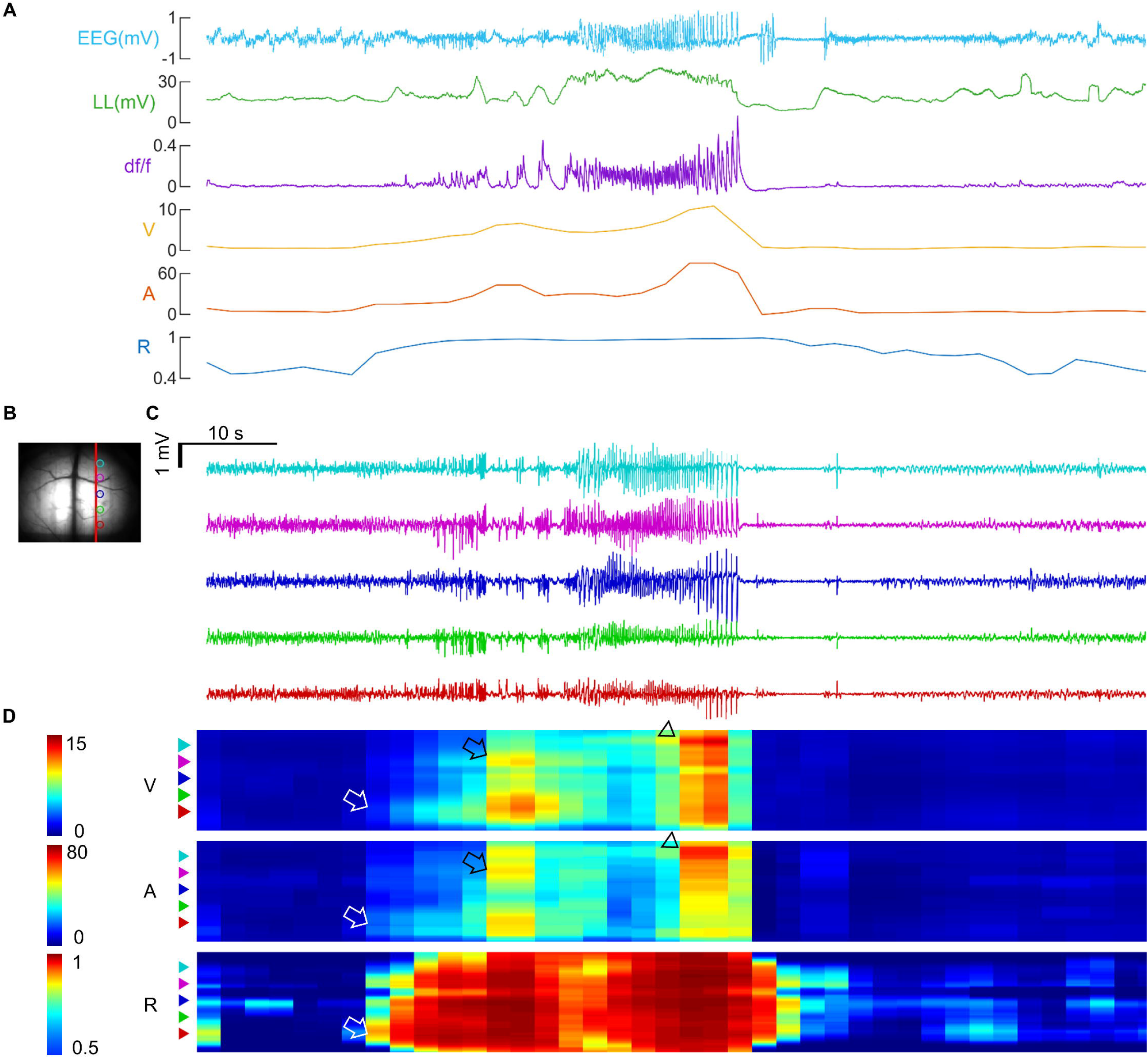
A representative seizure episode induced by chemogenetic suppression of cortical glutamatergic neurons. (A) From top to bottom: electrocorticogram (cyan), line length (LL, green), calcium signal (ΔF/F, purple), temporal variance (V, yellow), peak amplitude (A, orange), and synchrony ratio (R, red) over the course of the seizure. (B) Left: Raw fluorescence image of the dorsal cortex. The red vertical line indicates the cortical region along which the five ECoG channels were recorded, corresponding to the five traces in (C). (C) Representative ECoG traces from the five cortical sites, color-coded to match the locations in (B). (D) Spatiotemporal maps of V, A, and R derived from the red line in (B). The horizontal axis denotes time, identical to (A) and (C); the vertical axis corresponds to pixel positions along the red line, from top to bottom. The colored triangular markers on the left indicate the spatial positions of each ECoG channel, with colors matching (B) and (C). White arrows highlight the initial posterior parietal phase, black arrows mark the subsequent spread to frontal cortex, and black triangles denote the later frontal-dominant phase.

During this seizure, ECoG activity, line length (LL), and calcium fluorescence rose in parallel, accompanied by hemisphere-wide increases in temporal variance (*V*), peak amplitude (*A*), and synchrony ratio (*R*), marking the transition from baseline to highly synchronized, high-amplitude network activity (Figure 5A). Combined electrophysiology and calcium imaging (Figure 5B–D) revealed a biphasic evolution: initial epileptiform discharges with abnormal, high-amplitude oscillatory calcium signals confined to the posterior parietal region (white arrows, Figure 5D), followed by a spatially non-contiguous shift to a frontal-dominant phase (black arrows/triangles). Supplementary Video 1 provides higher spatiotemporal resolution, showing focal onset at the occipito-parietal-temporal junction near the lateral margin of the injection site (13–17 s), rapid establishment of a secondary focus in medial frontal cortex (≈33 s), and subsequent bilateral generalization. Concurrent behavioral video confirmed clinical correlates: focal clonic movements of the contralateral limb evolving into generalized tonic–clonic activity, consistent with the localized cortical origin evident in the combined ECoG–imaging data.

## Discussion

In this study, we report an unconventional finding: chemogenetic suppression of cortical glutamatergic neurons via Gi-coupled DREADDs paradoxically elicits focal interictal-like spikes and seizures, challenging the classical excitation–inhibition (E/I) model of epilepsy. By combining wide-field calcium imaging with high-density electrocorticography, we demonstrate that this paradoxical activity is reproducible in both transgenic and wild-type C57BL/6J mice, and that it generalizes across the dorsal cortex, appearing in parietal as well as frontal regions rather than being confined to a single locus. Importantly, some events progressed to overt seizures with clear behavioral correlates, confirming the epileptic nature of these discharges.

Our findings challenge the classical view that seizures arise only from excessive excitation or insufficient inhibition. Earlier slice work showed that tightly synchronised interneuronal bursts can precipitate interictal and ictal events by raising extracellular K and shifting the GABA-A receptor reversal potential ^13,14,16,17,36^. Here we demonstrate a complementary—and equally paradoxical—mechanism: chemogenetic suppression of glutamatergic neurons, without directly manipulating interneurons, can also reliably trigger focal epileptiform activity and, in some animals, full seizures in vivo.

At the single-cell level, hM4Di activation initiates Gi signaling that lowers neuronal excitability through several downstream actions. Giα suppresses adenylyl cyclase, reducing cAMP and thereby PKA and Epac activity. In neurons, this cascade has been shown to regulate multiple ion channels and receptors, including AMPA receptor downregulation ^37,38^, and upregulation of Kv4.2 potassium channels ^39^. In parallel, Giβγ activates GIRK channels, producing membrane hyperpolarization ^40,41^. Overall, these pathways are classically considered inhibitory. However, some downstream effectors can exert more complex effects on neuronal electrical properties. For example, lowering cAMP reduces HCN channel opening probability and shifts their voltage dependence ^42^, which increase input resistance, prolong membrane time constants, and enhance temporal summation — conditions that can favor synchronous activity. Clinically, HCN1 mutations have been associated with developmental epileptic encephalopathy ^43–45^ and HCN1 knockout mice have been demonstrated with increased seizure susceptibility ^46^, underscoring that, although paradoxical, cAMP–HCN modulation can contribute to pro-epileptic states. It is therefore conceivable that changes in neuronal electrical properties, even in hyperpolarized cells, might promote synchronization under some conditions.

Given that the dominant single-cell effect of hM4Di is suppression, the ictogenic mechanism is most plausibly circuit-based. Silencing a subset of excitatory neurons could reduce excitatory drive to local interneurons. PV interneurons, which are central to inhibitory control, are especially dependent on convergent and synchronous excitation (Hu et al., 2014), as their dendrites are electrically leaky ^47,48^ and excitatory synapses onto them are dominated by short-term depression ^49,50^. Thus, if part of the excitatory population is silenced, it is conceivable that PV interneurons may no longer receive sufficient convergent drive, diminishing their inhibitory output and allowing neighboring untransfected excitatory neurons to develop pathological activity. Consistent with this possibility, seizures in our model often initiated at the edge of the transfected zone (Supplementary Video 1), where suppressed and unsuppressed excitatory populations interface. Interestingly, theoretical work on balanced E/I networks has also shown that modest reductions in excitatory drive, in certain parameter regimes, can destabilize the asynchronous-irregular state and generate large, population-synchronous oscillations ^51,52^. These models do not incorporate the coincidence-detection properties of PV interneurons, yet they predict that reducing excitatory activity alone can paradoxically promote hypersynchrony. Our findings may represent one biological pathway through which such theoretical instabilities manifest in vivo.

Over longer timescales, sustained silencing can engage homeostatic plasticity mechanisms. Neuronal intrinsic excitability can rebound within minutes of reduced activity ^53,54^. If a large population of neighboring excitatory neurons undergoes such compensatory potentiation, the network may become hypersensitive to synchronizing inputs, potentially facilitating epileptiform activity. Synaptic rescaling of excitatory inputs can also occur when presynaptic drive is reduced, although this process typically unfolds over a longer timescale ^55^. By globally increasing postsynaptic gain, scaling can destabilize network balance and lower the threshold for runaway activity, creating conditions permissive for epileptiform discharges. Taken together, the paradoxical epileptiform activity induced by inhibitory DREADD activation is unlikely to arise from a single process. Instead, it may reflect the interaction of rapid circuit-level disfacilitation, network instabilities that emerge with reduced excitation, and slower homeostatic adaptations that amplify susceptibility over time.

Our findings have several clinically relevant implications. First, our data may help explain structural epilepsies associated with traumatic brain injury or stroke-induced cortical damage, conditions often characterized by substantial loss or dysfunction of glutamatergic neurons ^22,23^. Second, they highlight that therapeutic strategies which tip networks toward inhibition— whether by suppressing excitation, as in our model, or by pharmacologically enhancing GABAergic transmission—can paradoxically promote seizures in certain contexts. Indeed, vigabatrin and tiagabine, despite augmenting inhibition, have been reported to aggravate absence seizures or provoke seizure exacerbation in specific epilepsy syndromes ^56^. Finally, although our experiments do not model sleep physiology, the finding that seizures can emerge when excitation is suppressed parallels the clinical observation that seizures sometimes arise during inhibitory-dominant states such as NREM sleep ^18^. Together, these observations underscore the importance of nuanced epilepsy management strategies that account for cell-type, timing, and specific circuit dynamics, rather than uniformly enhancing inhibition. Achieving these clinical goals will require future mechanistic studies to delineate and validate the specific cellular and network processes underlying paradoxical instability. In parallel, identifying biomarkers or electrophysiological signatures in patients whose seizures arise from similar inhibition-driven mechanisms will be essential for guiding personalized therapeutic approaches.

## Acknowledgement

We acknowledge Mingrui Zhao for technical assistance, and Zhuohao Wu, Pearly Ye, and Wei Wang for experimental support.

## Funding

JYL, QWZ, and BG acknowledge funding from William Rhodes Center for Glioblastoma -Collaborative Research Initiative to accomplish this work. This material is based upon work supported by, or in part by, the Foundation for Anesthesia Education and Research.

### Supplementary material

Supplementary material is available at Brain online.

## Abbreviations

AAV: adeno-associated virus
CNO: clozapine-N-oxide
DREADD: Designer Receptor Exclusively Activated by Designer Drugs
hM3Dq: human muscarinic M3 receptor–derived excitatory DREADD
hM4Di: human muscarinic M4 receptor–derived inhibitory DREADD
IED: interictal epileptiform discharge

