## Supplementary Materials for "Suppressing cortical glutamatergic neurons produces paradoxical interictal discharges and seizures"

Supplementary Data 1: Viral Genome Sequencing Report

This document contains the sequencing validation data for the AAV9-CaMKIIα-hM3Dq-mCherry and AAV9-CaMKIIα-hM4Di-mCherry viral constructs. It includes summary reports, gel images, read distribution histograms, FASTA assemblies, and GenBank annotations.

### AAV9-CaMKIIα-hM3Dq-mCherry

#### Summary Report

1-mer (%) 2-mer (%) 3-mer (%)

moles	99.8	0.0	0.2

mass	99.5	0.0	0.5

*************************

E. coli genomic contamination: 0.0%

#### Gel Image

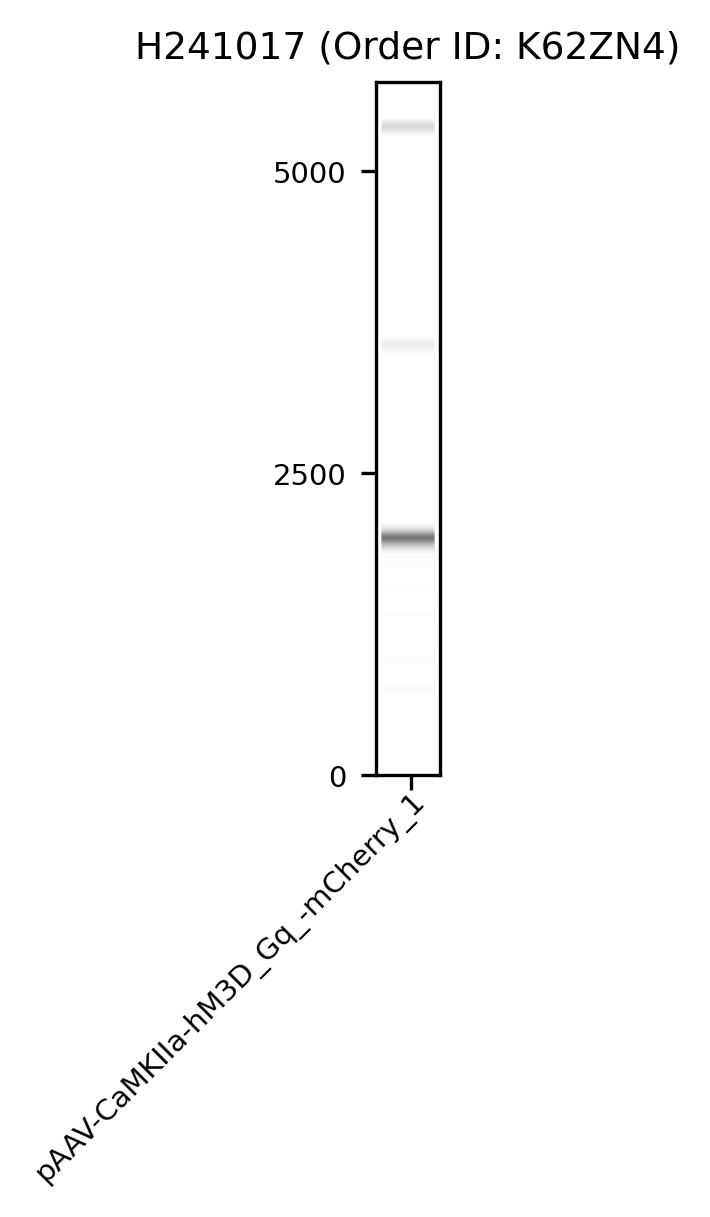

#### Histogram

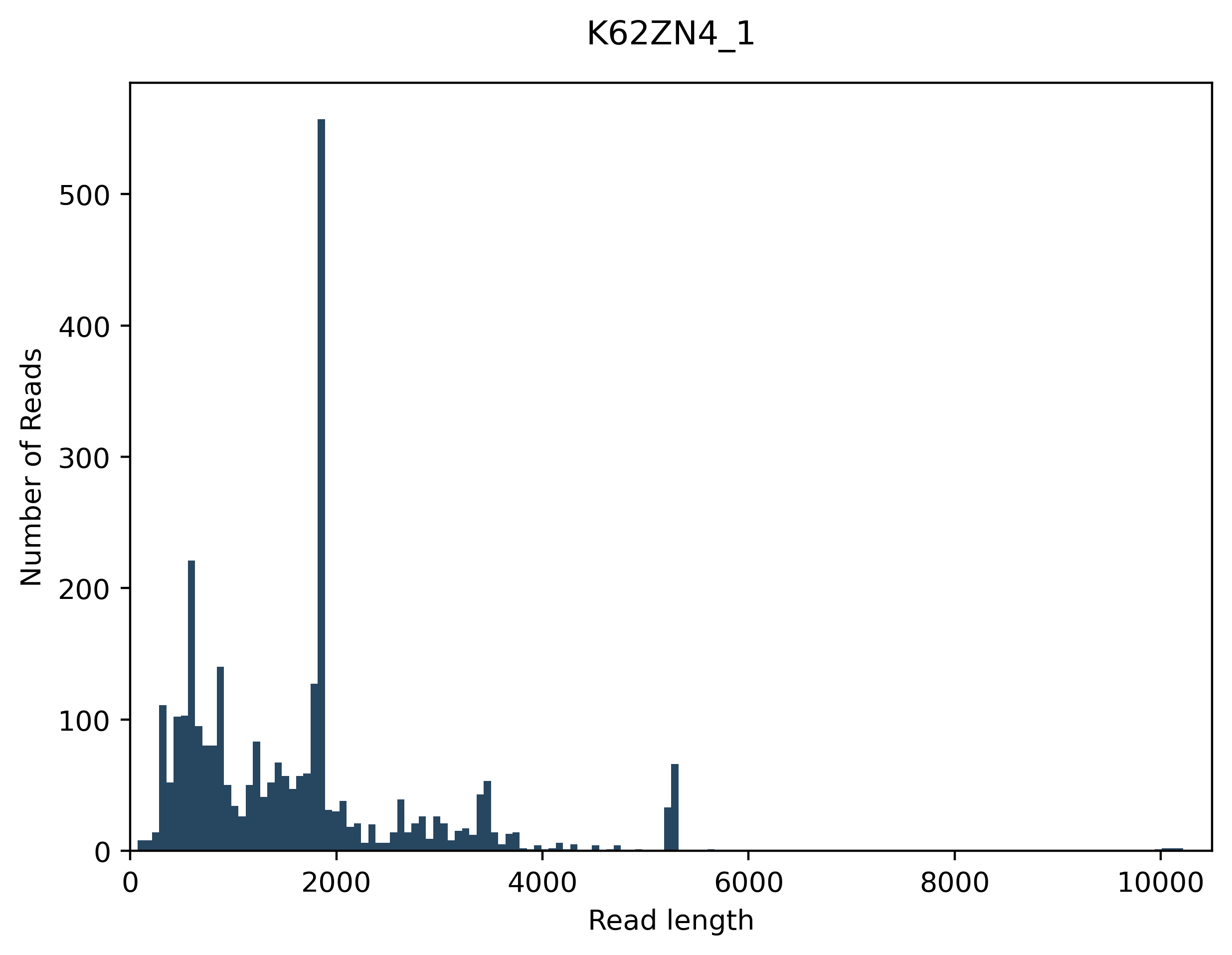

#### FASTA Assemblies

File: K62ZN4_1_assembly-1_pAAV-CaMKIIa-hM3D_Gq_-mCherry.fasta

>pAAV-CaMKIIa-hM3D_Gq_-mCherry

TTGGCCACTCCCTCTCTGCGCGCTCGCTCGCTCACTGAGGCCGCCCGGGCAAAGCCCGGG

CGTCGGGCGACCTTTGCTCGCGCGGCCTCAGTGAGCGAGCGAGCGCGCAGAGAGGGAGTG

GCCAACTCCATCACTAGGGGTTCCTGCGGCCGCTCGGTCCGCACGTGGTTACCTACAAAA

TCAGAAGGACAGGGAAGGGAGCAGTGGTTCACGCCTGTAATCCCAGCAATTTGGGAGGCC

AAGGTGGGTAGATCACCTGAGATTAGGAGTTGGAGACCAGCCTGGCCAATATGGTGAAAC

CCCGTCTCTACCAAAAAAACAAAAATTAGCTGAGCCTGGTCATGCATGCCTGGAATCCCA

ACAACTCGGGAGGCTGAGGCAGGAGAATCGCTTGAACCCAGGAGGCGGAGATTGCAGTGA

GCCAAGATTGTGCCACTGCACTCCAGCTTGGTTCCCAATAGACCCCGCAGGCCCTACAGG

TTGTCTTCCCAACTTGCCCCTTGCTCCATACCACCCCCCTCCACCCCATAATATTATAGA

AGGACACCTAGTCAGACAAAATGATGCAACTTAATTTTATTAGGACAAGGCTGGTGGGCA

CTGGAGTGGCAACTTCCAGGGCCAGGAGAGGCACTGGGGAGGGGTCACAGGGATGCCACC

CGTAGATCTCTCGAGCAGCGCTCGGTATCGATGCGGGGAGGCGGCCCAAAGGGAGATCCG

ACTCGTCTGAGGGCGAAGGCGAAGACGCGGAAGAGGCCGCAGAGCCGGCAGCAGGCCGCG

GGAAGGAAGGTCCGCTGGATTGAGGGCCGAAGGGACGTAGCAGAAGGACGTCCCGCGCAG

AATCCAGGTGGCAACATAGGCGAGCAGCCAAGGAAAGGACGATGATTTCCCCGACAACAC

CACGGCGGAACTCATCGCCGCCTGCCTTGCCCGCTGCTGGACAGGGGCTCGGCTGTTGGG

CACTGACAATTCCGTGGTGTTGTCGGGGAAATCATCGTCCTTTCCTTGGCTGCTCGCCTA

TGTTGCCACCTGGATTCTGCGCGGGACGTCCTTCTGCTACGTCCCTTCGGCCCTCAATCC

AGCGGACCTTCCTTCCCGCGGCCTGCTGCCGGCTCTGCGGCCTCTTCCGCGTCTTCGCCT

TCGCCCTCAGACGAGTCGGATCTCCCTTTGGGCCGCCTCCCCGCATCGATACCGAGCGCT

GCTCGAGAGATCTACGGGTGGCATCCCTGTGACCCCTCCCCAGTGCCTCTCCTGGCCCTG

GAAGTTGCCACTCCAGTGCCCACCAGCCTTGTCCTAATAAAATTAAGTTGCATCATTTTG

TCTGACTAGGTGTCCTTCTATAATATTATGGGGTGGAGGGGGGTGGTATGGAGCAAGGGG

CAAGTTGGGAAGACAACCTGTAGGGCCTGCGGGGTCTATTGGGAACCAAGCTGGAGTGCA

GTGGCACAATCTTGGCTCACTGCAATCTCCGCCTCCTGGGTTCAAGCGATTCTCCTGCCT

CAGCCTCCCGAGTTGTTGGGATTCCAGGCATGCATGACCAGGCTCAGCTAATTTTTGTTT

TTTTGGTAGAGACGGGGTTTCACCATATTGGCCAGGCTGGTCTCCAACTCCTAATCTCAG

GTGATCTACCCACCTTGGCCTCCCAAATTGCTGGGATTACAGGCGTGAACCACTGCTCCC

TTCCCTGTCCTTCTGATTTTGTAGGTAACCACGTGCGGACCGAGCGGCCGCAGGAACCCC

TAGTGATGGAGTTGGCCACTCCCTCTCTGCGCGCTCGCTCGCTCACTGAGGCCGCCCGGG

CAAAGGTCCCGGGCGTCGGGCGACCTTTGGTCGCCCGGCCTCAGTGAGCGAGCGAGCGCG

CAGAGAGGGAGTGGCCAAA

File: K62ZN4_1_assembly-2_pAAV-CaMKIIa-hM3D_Gq_-mCherry.fasta

>pAAV-CaMKIIa-hM3D_Gq_-mCherry

TTTGGCCACTCCCTCTCTGCGCGCTCGCTCGCTCACTGAGGCCGCCCGGGCAAAGCCCGG

GCGTCGGGCGACCTTTGGTCGCCCGGCCTCAGTGAGCGAGCGAGCGCGCAGAGAGGGAGT

GGCCAACTCCATCACTAGGGGTTCCTGCGGCCGCTCGGTCCGCACGTGGTTACCTACAAA

ATCAGAAGGACAGGGAAGGGAGCAGTGGTTCACGCCTGTAATCCCAGCAATTTGGGAGGC

CAAGGTGGGTAGATCACCTGAGATTAGGAGTTGGAGACCAGCCTGGCCAATATGGTGAAA

CCCCGTCTCTACCAAAAAAACAAAAATTAGCTGAGCCTGGTCATGCATGCCTGGAATCCC

AACAACTCGGGAGGCTGAGGCAGGAGAATCGCTTGAACCCAGGAGGCGGAGATTGCAGTG

AGCCAAGATTGTGCCACTGCACTCCAGCTTGGTTCCCAATAGACCCCGCAGGCCCTACAG

GTTGTCTTCCCAACTTGCCCCTTGCTCCATACCACCCCCCTCCACCCCATAATATTATAG

AAGGACACCTAGTCAGACAAAATGATGCAACTTAATTTTATTAGGACAAGGCTGGTGGGC

ACTGGAGTGGCAACTTCCAGGGCCAGGAGAGGCACTGGGGAGGGGTCACAGGGATGCCAC

CCGTAGATCTCTCGAGCAGCGCTCGGTATCGATGCGGGGAGGCGGCCCAAAGGGAGATCC

GACTCGTCTGAGGGCGAAGGCGAAGACGCGGAAGAGGCCGCAGAGCCGGCAGCAGGCCGC

GGGAAGGAAGGTCCGCTGGATTGAGGGCCGAAGGGACGTAGCAGAAGGACGTCCCGCGCA

GAATCCAGGTGGCAACATAGGCGAGCAGCCAAGGAAAGGACGATGATTTCCCCGACAACA

CCACGGAATTGTCAGTGCCCAACAGCCGAGCCCCTGTCCAGCAGCGGGCAAGGCAGGCGG

CGATGAGTTCCGCCGTGGCAATAGGGAGGGGGAAAGCGAAAGTTCCGGAAAGGAGCTGAC

AGGTGGTGGCAATGCCCCAACCAGTGGGGGTTGCGTCAGCAAACACAGTGCACACCACGC

CACGTTGCCTGACAACGGGCCACAACTCCTCATAAAGAGACAGCAACCAGGATTTATACA

AGGAGGAGAAAATGAAAGCCATACGGGAAGCAATAGCATGATACAAAGGCATTAAAGCAG

CGTATCCACATAGCGTAAAAGGAGCAACATAGTTAAGAATACCAGTCAATCTTTCACAAA

TTTTGTAATCCAGAGGTTGATTATCGATAAGCTTGATATCGAATTCTTACTTGTACAGCT

CGTCCATGCCGCCGGTGGAGTGGCGGCCCTCGGCGCGTTCGTACTGTTCCACGATGGTGT

AGTCCTCGTTGTGGGAGGTGATGTCCAACTTGATGTTGACGTTGTAGGCGCCGGGCAGCT

GCACGGGCTTCTTGGCCTTGTAGGTGGTCTTGACCTCAGCGTCGTAGTGGCCGCCGTCCT

TCAGCTTCAGCCTCTGCTTGATCTCGCCCTTCAGGGCGCCGTCCTCGGGGTACATCCGCT

CGGAGGAGGCCTCCCAGCCCATGGTCTTCTTCTGCATTACGGGGCCGTCGGAGGGGAAGT

TGGTGCCGCGCAGCTTCACCTTGTAGATGAACTCGCCGTCCTGCAGGGAGGAGTCCTGGG

TCACGGTCACCACGCCGCCGTCCTCGAAGTTCATCACGCGCTCCCACTTGAAGCCCTCGG

GGAAGGACAGCTTCAAGTAGTCGGGGATGTCGGCGGGGTGCTTCACGTAGGCCTTGGAGC

CGTACATGAACTGAGGGGACAGGATGTCCCAGGCGAAGGGCAGGGGGCCACCCTTGGTCA

CCTTCAGCTTGGCGGTCTGGGTGCCCTCGTAGGGGCGGCCCTCGCCCTCGCCCTCGATCT

CGAACTCGTGGCCGTTCACGGAGCCCTCCATGTGCACCTTGAAGCGCATGAACTCCTTGA

TGATGGCCATGTTATCCTCCTCGCCCTTGCTCACCATGGTGGCGACCGGGGGATCCTTCA

AGGCCTGCTCGGGTGCGCGCTTGTGAAAAATGACCGACTGTCTCTGCTGGTACTGCTGCT

TGCGCCTCTTTTTTTTGTCACACTGGCACAGCAGCAGCATCTTGAAAGTGGTTCTGAATG

TTTTGTTGCACAGAGCATAGCACACGGGGTTCACGGTGCTGTTGATGTAGCACAGCCAGT

AGCCCAGATTCCAAAAGGTTTTGGGTATGCAGCTGTCACAAAAGGTGTTCACCAGAACCA

TGATGTTGTATGGGGTCCAAGTGATGATGAAGGCAAGCAAGATCGCACTGAGGGTCTGGG

CCGCTTTCTTCTCCTTGACCAGGGACATCCTTTTCCGCTTAGTGATCTGACTTCTGGTCT

TCAGAGCAAACCTCTTGGCCAGAGTGGCTTCCTTGAAGGACAGAGGTAGAGTGGCCGTGC

TCTTACCCACTGAGGAGTTGACGTCAGAAGTCTTAGCTGTGTCCACGGCTGACTCTAGCT

GGATGGGAAGCTTGGAGAAGCTTTTTGGAAAACTGCCTCCATCGTCCACGCTCTTCTGGG

CCTGCAGCTTGTCGGCTTTCCTCTCCAAGTCCACCATCCCCAGCTCCTCCTCAGGCACCT

GCAGGTTGTCCGATGAGGGTAACTTGGTGGAGTTGAGGATGGTGCTGTGACCCGGAAGCT

TGAGCACGATGGAGTAGATGGCTCTCGTCTCGGAGCCAATGTCCTCCTCGTCGGAGGAGG

CGGAGTTCTCCAGGGAGGCAGCAGCATCATTGTTGTTCCAACTGTCACTGCTGCTGTGGT

CTTGGTCCATCTGCTCGGAGCTGGGTTTCCAGCTCTTGGTTGTGAACCAGAAGTGGCAGC

GGCCATACTTCCTCCTGTTGGAGCGTTTCATGCTTTGCTGTTGAAGTTCGTAACTGCTGC

AGCTTCGAGAACTGCCCGTGGGGTGGACAAAGTTTTCTGTCTCTGCCTCTGTCCCAGAGG

CTTGCAGGCCAGCAAGCTCTTTGGTACGCTTTTCAGTTTCCTTATAGATCCTCCAGTATA

AAATAGTCATAATGGTGACAGGCATATAAAAACCAGCGATGGCTGTGCCAAAAGTAATGG

TGGGCTCACTGAGGAACTGAATGAAGCACTCTCCCGGAGGCACAGTTCTCTTTCCAACAA

AGTATTGCCAGAACAAGATGGCAGGAGCCCAAAGGACAAAGGAGATGACCCAAGCCAGAC

CGATCATCACACCGGCTCTCTTTGTTGTTCGTTTGGCTCGGTACGTGAGCGGCCTCGTGA

TGGAAAAGTATCTGTCAAAGCTGATGACCAGAAGATTCATAACAGAGGCATTGCTGGCTA

CGCAGTCAATGGCAAGCCAGAGGTCACAGGCCAAGTTCCCTAAGGCCCATCGATTCATGA

TGATGTAGGTCGTAAACAGATTCATTGAAATGACCCCGATAATCAGATCGGCACAGGCCA

GGCTTAAGAGGAAGTAGTTGTTGACCGTCTTCAGCTGCTTGTTGACCTTAAATGACACAA

TTACCAGGATGTTGCCGATGATGGTCACCAAGGCCAGGATGCCCGTTAAGAAAGCGATGA

AGACCACTTGCCAGACGGTATGACCTCCCAGAGGGTCATCGGTGGTACCGTCTGGAGAGG

AGAAATTGCCAGCTGCTCGAGAAACATTGTAGCTGCCGAAATGAGTGACGGTTCCCGGGG

GCAGCCCTGCATCGGAGGGGCTGTGTATCCAGGAGGAGCTGATGTTTGGAAACAAAGGCG

AGGTTGTACTGTTATTGTGCAAGGTCATGGTGGCGTCGACTCTAGAGGATCCGGTACCGC

TCTAGAGCTGCCCCCAGAACTAGGGGCCACTCGCCTGCCCGTGCTCCTGAGTGCAAACGG

AGAACCGGCTTGACTGACGAGCTTGGGGCTTCTGAGCAGGGCACTGTGGCTGCTCACAGC

CTCGCGAGCCTTCGTCAGCATCCAGGTCCCCATGGCAACCACCTCCGAAACGCCCTGTTC

CCGTTGCCCCGGTAACTGCCTCCCCAACACCTGCCTGCCTTTCACTTTAAAACCTGCTCC

TCTTTGCCCTGGCCTTATATACACAATACTGCAGGACTGTGTGGGCCCAGGAGCAAGTGG

ACCCTGTTCCCCCAGGGTGATGTACTGAGGGGGTTGGAGAAGGGGATGCGAACAAACTTA

GTCCACAAGTGATCATCGCTCTTCCTACCTCCCCCACCTCCAACTATAGGGCCCTTTTAA

CAGACGGGATGGGATTAAGTGGGGCAAAGAGGCAGTTGCTATGGTAACGGCTAGTGGGTG

CCACATTCTGGGATGGTCCTTGAGAAAGACAGGATGTAGTTACCCTGGCAACTTCATCTC

CTCTGAAGAGACAGGTATTGCCCCGTTGCCCTGGCAACAGCTGGGTTCTAAGGTTTACAG

GGGAAGCTAATTCATCTCTCCTTCCCTTCCCCCTTGTAGTCTCTGTAATGGTTTCTTTGC

TTCCTCTGGGAATTGGTCTGGTTGACTTGGCCTCTGTCTCTGTCCTTGTTTCTTTCCAGC

TCTCCCTGCTTTTTAGCCATTGTGCATATCACACCGCCACCTTAGCGTGAGAAGAAGTAC

CAAACAGACCCAGAATGCTGACGGAATGACTCTGTCTGCTTTCTCACCCCTATCTGTCAC

CACTGAGCAGCAGGGCCCCTCCTGAGGCCTCTGCCACAGCTCTCCCCACCCATCCTCCCA

GAGGGACCATGAAGATTTGGTCTCCTGGGGCTCCTGTCCCAGTTTCTGCTTGTGCTTCCC

GTCAAGGTAGGACTGGAGAGAGGCAACTATAATGAGGTGAACGTGGCTGGACTCACCCAG

GGCTGGGCGGGTGGTGTAGATGTACCCTGCCCTGGACTCTTCTTTCCTAAACTCATTGGT

CATCTGCCCTAATGAGTTCACTCCTTCTGTATATCGTACTTGCATGCCCCCAAGGCATGG

AATTTAAGATTAGTAATCTGAGTTCAAGTTCTAGCTCTGACATTTCTGATGTGACCCTGG

GCAGGTCACTGAGGAAGCTCTCTGCACCCCCATCTCATTTTCTAGAAAATCAGAAGAAGA

ACCCCATGGAGATGAAGTGACCTAAGGCCATAATGTTAAACGCGTGCGGCCGCAGGAACC

CCTAGTGATGGAGTTGGCCACTCCCTCTCTGCGCGCTCGCTCGCTCACTGAGGCCGCGCG

ACCAAAGGTCGCCCGGCGTCGGGGACCTTTGCTCGGGCGGCCTCAGTGAGCGAGCGAGCG

CGCAGAGAGGGA

File: K62ZN4_1_assembly-3_pAAV-CaMKIIa-hM3D_Gq_-mCherry.fasta

pAAV-CaMKIIa-hM3D_Gq_-mCherry

TTGGCCACTCCCTCTCTGCGCGCTCGCTCGCTCACTGAGGCCGGGCGACCAAAGGTCGCC

CGACGTCGGGGACTTTGCTCGCGCGGCCTCAGTGAGCGAGCGAGCGCGCAGAGAGGGAGT

GGCCAACTCCATCACTAGGGGTTCCTGCGGCCGCTCGGTCCGCACGTGGTTACCTACAAA

ATCAGAAGGACAGGGAAGGGAGCAGTGGTTCACGCCTGTAATCCCAGCAATTTGGGAGGC

CAAGGTGGGTAGATCACCTGAGATTAGGAGTTGGAGACCAGCCTGGCCAATATGGTGAAA

CCCCGTCTCTACCAAAAAAACAAAAATTAGCTGAGCCTGGTCATGCATGCCTGGAATCCC

AACAACTCGGGAGGCTGAGGCAGGAGAATCGCTTGAACCCAGGAGGCGGAGATTGCAGTG

AGCCAAGATTGTGCCACTGCACTCCAGCTTGGTTCCCAATAGACCCCGCAGGCCCTACAG

GTTGTCTTCCCAACTTGCCCCTTGCTCCATACCACCCCCCTCCACCCCATAATATTATAG

AAGGACACCTAGTCAGACAAAATGATGCAACTTAATTTTATTAGGACAAGGCTGGTGGGC

ACTGGAGTGGCAACTTCCAGGGCCAGGAGAGGCACTGGGGAGGGGTCACAGGGATGCCAC

CCGTAGATCTCTCGAGCAGCGCTCGGTATCGATGCGGGGAGGCGGCCCAAAGGGAGATCC

GACTCGTCTGAGGGCGAAGGCGAAGACGCGGAAGAGGCCGCAGAGCCGGCAGCAGGCCGC

GGGAAGGAAGGTCCGCTGGATTGAGGGCCGAAGGGACGTAGCAGAAGGACGTCCCGCGCA

GAATCCAGGTGGCAACATAGGCGAGCAGCCAAGGAAAGGACGATGATTTCCCCGACAACA

CCACGGAATTGTCAGTGCCCAACAGCCGAGCCCCTGTCCAGCAGCGGGCAAGGCAGGCGG

CGATGAGTTCCGCCGTGGCAATAGGGAGGGGGAAAGCGAAAGTTCCGGAAAGGAGCTGAC

AGGTGGTGGCAATGCCCCAACCAGTGGGGGTTGCGTCAGCAAACACAGTGCACACCACGC

CACGTTGCCTGACAACGGGCCACAACTCCTCATAAAGAGACAGCAACCAGGATTTATACA

AGGAGGAGAAAATGAAAGCCATACGGGAAGCAATAGCATGATACAAAGGCATTAAAGCAG

CGTATCCACATAGCGTAAAAGGAGCAACATAGTTAAGAATACCAGTCAATCTTTCACAAA

TTTTGTAATCCAGAGGTTGATTATCGATAAGCTTGATATCGAATTCTTACTTGTACAGCT

CGTCCATGCCGCCGGTGGAGTGGCGGCCCTCGGCGCGTTCGTACTGTTCCACGATGGTGT

AGTCCTCGTTGTGGGAGGTGATGTCCAACTTGATGTTGACGTTGTAGGCGCCGGGCAGCT

GCACGGGCTTCTTGGCCTTGTAGGTGGTCTTGACCTCAGCGTCGTAGTGGCCGCCGTCCT

TCAGCTTCAGCCTCTGCTTGATCTCGCCCTTCAGGGCGCCGTCCTCGGGGTACATCCGCT

CGGAGGAGGCCTCCCAGCCCATGGTCTTCTTCTGCATTACGGGGCCGTCGGAGGGGAAGT

TGGTGCCGCGCAGCTTCACCTTGTAGATGAACTCGCCGTCCTGCAGGGAGGAGTCCTGGG

TCACGGTCACCACGCCGCCGTCCTCGAAGTTCATCACGCGCTCCCACTTGAAGCCCTCGG

GGAAGGACAGCTTCAAGTAGTCGGGGATGTCGGCGGGGTGCTTCACGTAGGCCTTGGAGC

CGTACATGAACTGAGGGGACGGCGGCGTGGTGACCGTGACCCAGGACTCCTCCCTGCAGG

ACGGCGAGTTCATCTACAAGGTGAAGCTGCGCGGCACCAACTTCCCCTCCGACGGCCCCG

TAATGCAGAAGAAGACCATGGGCTGGGAGGCCTCCTCCGAGCGGATGTACCCCGAGGACG

GCGCCCTGAAGGGCGAGATCAAGCAGAGGCTGAAGCTGAAGGACGGCGGCCACTACGACG

CTGAGGTCAAGACCACCTACAAGGCCAAGAAGCCCGTGCAGCTGCCCGGCGCCTACAACG

TCAACATCAAGTTGGACATCACCTCCCACAACGAGGACTACACCATCGTGGAACAGTACG

AACGCGCCGAGGGCCGCCACTCCACCGGCGGCATGGACGAGCTGTACAAGTAAGAATTCG

ATATCAAGCTTATCGATAATCAACCTCTGGATTACAAAATTTGTGAAAGATTGACTGGTA

TTCTTAACTATGTTGCTCCTTTTACGCTATGTGGATACGCTGCTTTAATGCCTTTGTATC

ATGCTATTGCTTCCCGTATGGCTTTCATTTTCTCCTCCTTGTATAAATCCTGGTTGCTGT

CTCTTTATGAGGAGTTGTGGCCCGTTGTCAGGCAACGTGGCGTGGTGTGCACTGTGTTTG

CTGACGCAACCCCCACTGGTTGGGGCATTGCCACCACCTGTCAGCTCCTTTCCGGAACTT

TCGCTTTCCCCCTCCCTATTGCCACGGCGGAACTCATCGCCGCCTGCCTTGCCCGCTGCT

GGACAGGGGCTCGGCTGTTGGGCACTGACAATTCCGTGGTGTTGTCGGGGAAATCATCGT

CCTTTCCTTGGCTGCTCGCCTATGTTGCCACCTGGATTCTGCGCGGGACGTCCTTCTGCT

ACGTCCCTTCGGCCCTCAATCCAGCGGACCTTCCTTCCCGCGGCCTGCTGCCGGCTCTGC

GGCCTCTTCCGCGTCTTCGCCTTCGCCCTCAGACGAGTCGGATCTCCCTTTGGGCCGCCT

CCCCGCATCGATACCGAGCGCTGCTCGAGAGATCTACGGGTGGCATCCCTGTGACCCCTC

CCCAGTGCCTCTCCTGGCCCTGGAAGTTGCCACTCCAGTGCCCACCAGCCTTGTCCTAAT

AAAATTAAGTTGCATCATTTTGTCTGACTAGGTGTCCTTCTATAATATTATGGGGTGGAG

GGGGGTGGTATGGAGCAAGGGGCAAGTTGGGAAGACAACCTGTAGGGCCTGCGGGGTCTA

TTGGGAACCAAGCTGGAGTGCAGTGGCACAATCTTGGCTCACTGCAATCTCCGCCTCCTG

GGTTCAAGCGATTCTCCTGCCTCAGCCTCCCGAGTTGTTGGGATTCCAGGCATGCATGAC

CAGGCTCAGCTAATTTTTGTTTTTTTGGTAGAGACGGGGTTTCACCATATTGGCCAGGCT

GGTCTCCAACTCCTAATCTCAGGTGATCTACCCACCTTGGCCTCCCAAATTGCTGGGATT

ACAGGCGTGAACCACTGCTCCCTTCCCTGTCCTTCTGATTTTGTAGGTAACCACGTGCGG

ACCGAGCGGCCGCAGGAACCCCTAGTGATGGAGTTGGCCACTCCCTCTCTGCGCGCTCGC

TCGCTCACTGAGGCCGGGCGACCAAAGGTCGCCCGACGCCGGGGCTTTGCCCGGGCGGCC

TCAGTGAGCGAGCGAGCGCGCAGAGAGGGAGTGGCCAAA

#### GenBank Annotations

File: K62ZN4_1_assembly-1_pAAV-CaMKIIa-hM3D_Gq_-mCherry.gbk

LOCUS	pAAV-CaMKIIa-hM3D_Gq_-mCherry 1879 bp	DNA	linear	SYN 26-OCT-2024

DEFINITION pAAV-CaMKIIa-hM3D_Gq_-mCherry.

ACCESSION	pAAV-CaMKIIa-hM3D_Gq_-mCherry

VERSION	pAAV-CaMKIIa-hM3D_Gq_-mCherry

KEYWORDS	.

SOURCE

ORGANISM .

.

COMMENT	Annotated with pLannotate v1.2.2

FEATURES	Location/Qualifiers

polyA_signal	1216..1692

/note="pLannotate"

/label="hGH poly(A) signal"

/database="snapgene"

/identity="100.0"

/match_length="100.0"

/fragment="False"

/other="polyA_signal"

polyA_signal	complement(185..661)

/note="pLannotate"

/label="hGH poly(A) signal"

/database="snapgene"

/identity="100.0"

/match_length="100.0"

/fragment="False"

/other="polyA_signal"

misc_feature	900..1184

/note="pLannotate"

/label="WPRE (fragment)"

/database="snapgene"

/identity="99.6"

/match_length="48.4"

/fragment="True"

/other="misc_feature"

repeat_region	complement(16..145)

/note="pLannotate"

/label="AAV2 ITR (fragment)"

/database="snapgene"

/identity="98.5"

/match_length="92.2"

/fragment="True"

/other="repeat_region"

repeat_region	complement(1767..1878)

/note="pLannotate"

/label="AAV2 ITR (fragment)"

/database="snapgene"

/identity="98.2"

/match_length="79.4"

/fragment="True"

/other="repeat_region"

misc_feature	complement(693..905)

/note="pLannotate"

/label="WPRE (fragment)"

/database="snapgene"

/identity="99.5"

/match_length="36.2"

/fragment="True"

/other="misc_feature"

ORIGIN

1 ttggccactc cctctctgcg cgctcgctcg ctcactgagg ccgcccgggc aaagcccggg

61 cgtcgggcga cctttgctcg cgcggcctca gtgagcgagc gagcgcgcag agagggagtg

121 gccaactcca tcactagggg ttcctgcggc cgctcggtcc gcacgtggtt acctacaaaa

181 tcagaaggac agggaaggga gcagtggttc acgcctgtaa tcccagcaat ttgggaggcc

241 aaggtgggta gatcacctga gattaggagt tggagaccag cctggccaat atggtgaaac

301 cccgtctcta ccaaaaaaac aaaaattagc tgagcctggt catgcatgcc tggaatccca

361 acaactcggg aggctgaggc aggagaatcg cttgaaccca ggaggcggag attgcagtga

421 gccaagattg tgccactgca ctccagcttg gttcccaata gaccccgcag gccctacagg

481 ttgtcttccc aacttgcccc ttgctccata ccacccccct ccaccccata atattataga

541 aggacaccta gtcagacaaa atgatgcaac ttaattttat taggacaagg ctggtgggca

601 ctggagtggc aacttccagg gccaggagag gcactgggga ggggtcacag ggatgccacc

661 cgtagatctc tcgagcagcg ctcggtatcg atgcggggag gcggcccaaa gggagatccg

721 actcgtctga gggcgaaggc gaagacgcgg aagaggccgc agagccggca gcaggccgcg

781 ggaaggaagg tccgctggat tgagggccga agggacgtag cagaaggacg tcccgcgcag

841 aatccaggtg gcaacatagg cgagcagcca aggaaaggac gatgatttcc ccgacaacac

901 cacggcggaa ctcatcgccg cctgccttgc ccgctgctgg acaggggctc ggctgttggg

961 cactgacaat tccgtggtgt tgtcggggaa atcatcgtcc tttccttggc tgctcgccta

1021 tgttgccacc tggattctgc gcgggacgtc cttctgctac gtcccttcgg ccctcaatcc

1081 agcggacctt ccttcccgcg gcctgctgcc ggctctgcgg cctcttccgc gtcttcgcct

1141 tcgccctcag acgagtcgga tctccctttg ggccgcctcc ccgcatcgat accgagcgct

1201 gctcgagaga tctacgggtg gcatccctgt gacccctccc cagtgcctct cctggccctg

1261 gaagttgcca ctccagtgcc caccagcctt gtcctaataa aattaagttg catcattttg

1321 tctgactagg tgtccttcta taatattatg gggtggaggg gggtggtatg gagcaagggg

1381 caagttggga agacaacctg tagggcctgc ggggtctatt gggaaccaag ctggagtgca

1441 gtggcacaat cttggctcac tgcaatctcc gcctcctggg ttcaagcgat tctcctgcct

1501 cagcctcccg agttgttggg attccaggca tgcatgacca ggctcagcta atttttgttt

1561 ttttggtaga gacggggttt caccatattg gccaggctgg tctccaactc ctaatctcag

1621 gtgatctacc caccttggcc tcccaaattg ctgggattac aggcgtgaac cactgctccc

1681 ttccctgtcc ttctgatttt gtaggtaacc acgtgcggac cgagcggccg caggaacccc

1741 tagtgatgga gttggccact ccctctctgc gcgctcgctc gctcactgag gccgcccggg

1801 caaaggtccc gggcgtcggg cgacctttgg tcgcccggcc tcagtgagcg agcgagcgcg

1861 cagagaggga gtggccaaa

//File: K62ZN4_1_assembly-2_pAAV-CaMKIIa-hM3D_Gq_-mCherry.gbk

LOCUS	pAAV-CaMKIIa-hM3D_Gq_-mCherry 5292 bp	DNA	linear	SYN 26-OCT-2024

DEFINITION pAAV-CaMKIIa-hM3D_Gq_-mCherry.

ACCESSION	pAAV-CaMKIIa-hM3D_Gq_-mCherry

VERSION	pAAV-CaMKIIa-hM3D_Gq_-mCherry

KEYWORDS	.

SOURCE

ORGANISM .

.

COMMENT	Annotated with pLannotate v1.2.2

FEATURES	Location/Qualifiers

promoter	complement(3847..5135)

/note="pLannotate"

/label="CaMKII promoter"

/database="snapgene"

/identity="100.0"

/match_length="100.0"

/fragment="False"

/other="promoter"

CDS	complement(1310..2017)

/note="pLannotate"

/label="mCherry"

/database="fpbase"

/identity="100.0"

/match_length="100.0"

/fragment="False"

/other="CDS"

polyA_signal	complement(186..662)

/note="pLannotate"

/label="hGH poly(A) signal"

/database="snapgene"

/identity="100.0"

/match_length="100.0"

/fragment="False"

/other="polyA_signal"

CDS	complement(2039..3808)

/note="pLannotate"

/label="CHRM3"

/database="swissprot"

/identity="99.7"

/match_length="100.0"

/fragment="False"

/other="CDS"

misc_feature	complement(694..1282)

/note="pLannotate"

/label="WPRE"

/database="snapgene"

/identity="99.7"

/match_length="100.0"

/fragment="False"

/other="misc_feature"

repeat_region	complement(17..146)

/note="pLannotate"

/label="AAV2 ITR (fragment)"

/database="snapgene"

/identity="100.0"

/match_length="92.2"

/fragment="True"

/other="repeat_region"

misc_feature	complement(1282..1306)

/note="pLannotate"

/label="MCS (fragment)"

/database="snapgene"

/identity="100.0"

/match_length="23.1"

/fragment="True"

/other="misc_feature"

misc_feature	complement(3815..3832)

/note="pLannotate"

/label="MCS (fragment)"

/database="snapgene"

/identity="100.0"

/match_length="31.6"

/fragment="True"

/other="misc_feature"

ORIGIN

1 tttggccact ccctctctgc gcgctcgctc gctcactgag gccgcccggg caaagcccgg

61 gcgtcgggcg acctttggtc gcccggcctc agtgagcgag cgagcgcgca gagagggagt

121 ggccaactcc atcactaggg gttcctgcgg ccgctcggtc cgcacgtggt tacctacaaa

181 atcagaagga cagggaaggg agcagtggtt cacgcctgta atcccagcaa tttgggaggc

241 caaggtgggt agatcacctg agattaggag ttggagacca gcctggccaa tatggtgaaa

301 ccccgtctct accaaaaaaa caaaaattag ctgagcctgg tcatgcatgc ctggaatccc

361 aacaactcgg gaggctgagg caggagaatc gcttgaaccc aggaggcgga gattgcagtg

421 agccaagatt gtgccactgc actccagctt ggttcccaat agaccccgca ggccctacag

481 gttgtcttcc caacttgccc cttgctccat accacccccc tccaccccat aatattatag

541 aaggacacct agtcagacaa aatgatgcaa cttaatttta ttaggacaag gctggtgggc

601 actggagtgg caacttccag ggccaggaga ggcactgggg aggggtcaca gggatgccac

661 ccgtagatct ctcgagcagc gctcggtatc gatgcgggga ggcggcccaa agggagatcc

721 gactcgtctg agggcgaagg cgaagacgcg gaagaggccg cagagccggc agcaggccgc

781 gggaaggaag gtccgctgga ttgagggccg aagggacgta gcagaaggac gtcccgcgca

841 gaatccaggt ggcaacatag gcgagcagcc aaggaaagga cgatgatttc cccgacaaca

901 ccacggaatt gtcagtgccc aacagccgag cccctgtcca gcagcgggca aggcaggcgg

961 cgatgagttc cgccgtggca atagggaggg ggaaagcgaa agttccggaa aggagctgac

1021 aggtggtggc aatgccccaa ccagtggggg ttgcgtcagc aaacacagtg cacaccacgc

1081 cacgttgcct gacaacgggc cacaactcct cataaagaga cagcaaccag gatttataca

1141 aggaggagaa aatgaaagcc atacgggaag caatagcatg atacaaaggc attaaagcag

1201 cgtatccaca tagcgtaaaa ggagcaacat agttaagaat accagtcaat ctttcacaaa

1261 ttttgtaatc cagaggttga ttatcgataa gcttgatatc gaattcttac ttgtacagct

1321 cgtccatgcc gccggtggag tggcggccct cggcgcgttc gtactgttcc acgatggtgt

1381 agtcctcgtt gtgggaggtg atgtccaact tgatgttgac gttgtaggcg ccgggcagct

1441 gcacgggctt cttggccttg taggtggtct tgacctcagc gtcgtagtgg ccgccgtcct

1501 tcagcttcag cctctgcttg atctcgccct tcagggcgcc gtcctcgggg tacatccgct

1561 cggaggaggc ctcccagccc atggtcttct tctgcattac ggggccgtcg gaggggaagt

1621 tggtgccgcg cagcttcacc ttgtagatga actcgccgtc ctgcagggag gagtcctggg

1681 tcacggtcac cacgccgccg tcctcgaagt tcatcacgcg ctcccacttg aagccctcgg

1741 ggaaggacag cttcaagtag tcggggatgt cggcggggtg cttcacgtag gccttggagc

1801 cgtacatgaa ctgaggggac aggatgtccc aggcgaaggg cagggggcca cccttggtca

1861 ccttcagctt ggcggtctgg gtgccctcgt aggggcggcc ctcgccctcg ccctcgatct

1921 cgaactcgtg gccgttcacg gagccctcca tgtgcacctt gaagcgcatg aactccttga

1981 tgatggccat gttatcctcc tcgcccttgc tcaccatggt ggcgaccggg ggatccttca

2041 aggcctgctc gggtgcgcgc ttgtgaaaaa tgaccgactg tctctgctgg tactgctgct

2101 tgcgcctctt ttttttgtca cactggcaca gcagcagcat cttgaaagtg gttctgaatg

2161 ttttgttgca cagagcatag cacacggggt tcacggtgct gttgatgtag cacagccagt

2221 agcccagatt ccaaaaggtt ttgggtatgc agctgtcaca aaaggtgttc accagaacca

2281 tgatgttgta tggggtccaa gtgatgatga aggcaagcaa gatcgcactg agggtctggg

2341 ccgctttctt ctccttgacc agggacatcc ttttccgctt agtgatctga cttctggtct

2401 tcagagcaaa cctcttggcc agagtggctt ccttgaagga cagaggtaga gtggccgtgc

2461 tcttacccac tgaggagttg acgtcagaag tcttagctgt gtccacggct gactctagct

2521 ggatgggaag cttggagaag ctttttggaa aactgcctcc atcgtccacg ctcttctggg

2581 cctgcagctt gtcggctttc ctctccaagt ccaccatccc cagctcctcc tcaggcacct

2641 gcaggttgtc cgatgagggt aacttggtgg agttgaggat ggtgctgtga cccggaagct

2701 tgagcacgat ggagtagatg gctctcgtct cggagccaat gtcctcctcg tcggaggagg

2761 cggagttctc cagggaggca gcagcatcat tgttgttcca actgtcactg ctgctgtggt

2821 cttggtccat ctgctcggag ctgggtttcc agctcttggt tgtgaaccag aagtggcagc

2881 ggccatactt cctcctgttg gagcgtttca tgctttgctg ttgaagttcg taactgctgc

2941 agcttcgaga actgcccgtg gggtggacaa agttttctgt ctctgcctct gtcccagagg

3001 cttgcaggcc agcaagctct ttggtacgct tttcagtttc cttatagatc ctccagtata

3061 aaatagtcat aatggtgaca ggcatataaa aaccagcgat ggctgtgcca aaagtaatgg

3121 tgggctcact gaggaactga atgaagcact ctcccggagg cacagttctc tttccaacaa

3181 agtattgcca gaacaagatg gcaggagccc aaaggacaaa ggagatgacc caagccagac

3241 cgatcatcac accggctctc tttgttgttc gtttggctcg gtacgtgagc ggcctcgtga

3301 tggaaaagta tctgtcaaag ctgatgacca gaagattcat aacagaggca ttgctggcta

3361 cgcagtcaat ggcaagccag aggtcacagg ccaagttccc taaggcccat cgattcatga

3421 tgatgtaggt cgtaaacaga ttcattgaaa tgaccccgat aatcagatcg gcacaggcca

3481 ggcttaagag gaagtagttg ttgaccgtct tcagctgctt gttgacctta aatgacacaa

3541 ttaccaggat gttgccgatg atggtcacca aggccaggat gcccgttaag aaagcgatga

3601 agaccacttg ccagacggta tgacctccca gagggtcatc ggtggtaccg tctggagagg

3661 agaaattgcc agctgctcga gaaacattgt agctgccgaa atgagtgacg gttcccgggg

3721 gcagccctgc atcggagggg ctgtgtatcc aggaggagct gatgtttgga aacaaaggcg

3781 aggttgtact gttattgtgc aaggtcatgg tggcgtcgac tctagaggat ccggtaccgc

3841 tctagagctg cccccagaac taggggccac tcgcctgccc gtgctcctga gtgcaaacgg

3901 agaaccggct tgactgacga gcttggggct tctgagcagg gcactgtggc tgctcacagc

3961 ctcgcgagcc ttcgtcagca tccaggtccc catggcaacc acctccgaaa cgccctgttc

4021 ccgttgcccc ggtaactgcc tccccaacac ctgcctgcct ttcactttaa aacctgctcc

4081 tctttgccct ggccttatat acacaatact gcaggactgt gtgggcccag gagcaagtgg

4141 accctgttcc cccagggtga tgtactgagg gggttggaga aggggatgcg aacaaactta

4201 gtccacaagt gatcatcgct cttcctacct cccccacctc caactatagg gcccttttaa

4261 cagacgggat gggattaagt ggggcaaaga ggcagttgct atggtaacgg ctagtgggtg

4321 ccacattctg ggatggtcct tgagaaagac aggatgtagt taccctggca acttcatctc

4381 ctctgaagag acaggtattg ccccgttgcc ctggcaacag ctgggttcta aggtttacag

4441 gggaagctaa ttcatctctc cttcccttcc cccttgtagt ctctgtaatg gtttctttgc

4501 ttcctctggg aattggtctg gttgacttgg cctctgtctc tgtccttgtt tctttccagc

4561 tctccctgct ttttagccat tgtgcatatc acaccgccac cttagcgtga gaagaagtac

4621 caaacagacc cagaatgctg acggaatgac tctgtctgct ttctcacccc tatctgtcac

4681 cactgagcag cagggcccct cctgaggcct ctgccacagc tctccccacc catcctccca

4741 gagggaccat gaagatttgg tctcctgggg ctcctgtccc agtttctgct tgtgcttccc

4801 gtcaaggtag gactggagag aggcaactat aatgaggtga acgtggctgg actcacccag

4861 ggctgggcgg gtggtgtaga tgtaccctgc cctggactct tctttcctaa actcattggt

4921 catctgccct aatgagttca ctccttctgt atatcgtact tgcatgcccc caaggcatgg

4981 aatttaagat tagtaatctg agttcaagtt ctagctctga catttctgat gtgaccctgg

5041 gcaggtcact gaggaagctc tctgcacccc catctcattt tctagaaaat cagaagaaga

5101 accccatgga gatgaagtga cctaaggcca taatgttaaa cgcgtgcggc cgcaggaacc

5161 cctagtgatg gagttggcca ctccctctct gcgcgctcgc tcgctcactg aggccgcgcg

5221 accaaaggtc gcccggcgtc ggggaccttt gctcgggcgg cctcagtgag cgagcgagcg

5281 cgcagagagg ga

//File: K62ZN4_1_assembly-3_pAAV-CaMKIIa-hM3D_Gq_-mCherry.gbk

LOCUS	pAAV-CaMKIIa-hM3D_Gq_-mCherry 3519 bp	DNA	linear	SYN 26-OCT-2024

DEFINITION pAAV-CaMKIIa-hM3D_Gq_-mCherry.

ACCESSION	pAAV-CaMKIIa-hM3D_Gq_-mCherry

VERSION	pAAV-CaMKIIa-hM3D_Gq_-mCherry

KEYWORDS	.

SOURCE

ORGANISM .

.

COMMENT	Annotated with pLannotate v1.2.2

FEATURES	Location/Qualifiers

polyA_signal	complement(186..662)

/note="pLannotate"

/label="hGH poly(A) signal"

/database="snapgene"

/identity="100.0"

/match_length="100.0"

/fragment="False"

/other="polyA_signal"

polyA_signal	2858..3334

/note="pLannotate"

/label="hGH poly(A) signal"

/database="snapgene"

/identity="100.0"

/match_length="100.0"

/fragment="False"

/other="polyA_signal"

misc_feature	complement(694..1282)

/note="pLannotate"

/label="WPRE"

/database="snapgene"

/identity="99.7"

/match_length="100.0"

/fragment="False"

/other="misc_feature"

misc_feature	2238..2826

/note="pLannotate"

/label="WPRE"

/database="snapgene"

/identity="99.7"

/match_length="100.0"

/fragment="False"

/other="misc_feature"

CDS	complement(1308..1820)

/note="pLannotate"

/label="mCherry (fragment)"

/database="snapgene"

/identity="100.0"

/match_length="72.2"

/fragment="True"

/other="CDS"

CDS	1806..2210

/note="pLannotate"

/label="Blue102 (fragment)"

/database="fpbase"

/identity="97.8"

/match_length="58.4"

/fragment="True"

/other="CDS"

repeat_region	3374..3503

/note="pLannotate"

/label="AAV2 ITR (fragment)"

/database="snapgene"

/identity="99.2"

/match_length="92.2"

/fragment="True"

/other="repeat_region"

repeat_region	1..111

/note="pLannotate"

/label="AAV2 ITR (fragment)"

/database="snapgene"

/identity="95.5"

/match_length="78.7"

/fragment="True"

/other="repeat_region"

misc_feature	2214..2238

/note="pLannotate"

/label="MCS (fragment)"

/database="snapgene"

/identity="100.0"

/match_length="23.1"

/fragment="True"

/other="misc_feature"

misc_feature	complement(1282..1306)

/note="pLannotate"

/label="MCS (fragment)"

/database="snapgene"

/identity="100.0"

/match_length="23.1"

/fragment="True"

/other="misc_feature"

ORIGIN

1 ttggccactc cctctctgcg cgctcgctcg ctcactgagg ccgggcgacc aaaggtcgcc

61 cgacgtcggg gactttgctc gcgcggcctc agtgagcgag cgagcgcgca gagagggagt

121 ggccaactcc atcactaggg gttcctgcgg ccgctcggtc cgcacgtggt tacctacaaa

181 atcagaagga cagggaaggg agcagtggtt cacgcctgta atcccagcaa tttgggaggc

241 caaggtgggt agatcacctg agattaggag ttggagacca gcctggccaa tatggtgaaa

301 ccccgtctct accaaaaaaa caaaaattag ctgagcctgg tcatgcatgc ctggaatccc

361 aacaactcgg gaggctgagg caggagaatc gcttgaaccc aggaggcgga gattgcagtg

421 agccaagatt gtgccactgc actccagctt ggttcccaat agaccccgca ggccctacag

481 gttgtcttcc caacttgccc cttgctccat accacccccc tccaccccat aatattatag

541 aaggacacct agtcagacaa aatgatgcaa cttaatttta ttaggacaag gctggtgggc

601 actggagtgg caacttccag ggccaggaga ggcactgggg aggggtcaca gggatgccac

661 ccgtagatct ctcgagcagc gctcggtatc gatgcgggga ggcggcccaa agggagatcc

721 gactcgtctg agggcgaagg cgaagacgcg gaagaggccg cagagccggc agcaggccgc

781 gggaaggaag gtccgctgga ttgagggccg aagggacgta gcagaaggac gtcccgcgca

841 gaatccaggt ggcaacatag gcgagcagcc aaggaaagga cgatgatttc cccgacaaca

901 ccacggaatt gtcagtgccc aacagccgag cccctgtcca gcagcgggca aggcaggcgg

961 cgatgagttc cgccgtggca atagggaggg ggaaagcgaa agttccggaa aggagctgac

1021 aggtggtggc aatgccccaa ccagtggggg ttgcgtcagc aaacacagtg cacaccacgc

1081 cacgttgcct gacaacgggc cacaactcct cataaagaga cagcaaccag gatttataca

1141 aggaggagaa aatgaaagcc atacgggaag caatagcatg atacaaaggc attaaagcag

1201 cgtatccaca tagcgtaaaa ggagcaacat agttaagaat accagtcaat ctttcacaaa

1261 ttttgtaatc cagaggttga ttatcgataa gcttgatatc gaattcttac ttgtacagct

1321 cgtccatgcc gccggtggag tggcggccct cggcgcgttc gtactgttcc acgatggtgt

1381 agtcctcgtt gtgggaggtg atgtccaact tgatgttgac gttgtaggcg ccgggcagct

1441 gcacgggctt cttggccttg taggtggtct tgacctcagc gtcgtagtgg ccgccgtcct

1501 tcagcttcag cctctgcttg atctcgccct tcagggcgcc gtcctcgggg tacatccgct

1561 cggaggaggc ctcccagccc atggtcttct tctgcattac ggggccgtcg gaggggaagt

1621 tggtgccgcg cagcttcacc ttgtagatga actcgccgtc ctgcagggag gagtcctggg

1681 tcacggtcac cacgccgccg tcctcgaagt tcatcacgcg ctcccacttg aagccctcgg

1741 ggaaggacag cttcaagtag tcggggatgt cggcggggtg cttcacgtag gccttggagc

1801 cgtacatgaa ctgaggggac ggcggcgtgg tgaccgtgac ccaggactcc tccctgcagg

1861 acggcgagtt catctacaag gtgaagctgc gcggcaccaa cttcccctcc gacggccccg

1921 taatgcagaa gaagaccatg ggctgggagg cctcctccga gcggatgtac cccgaggacg

1981 gcgccctgaa gggcgagatc aagcagaggc tgaagctgaa ggacggcggc cactacgacg

2041 ctgaggtcaa gaccacctac aaggccaaga agcccgtgca gctgcccggc gcctacaacg

2101 tcaacatcaa gttggacatc acctcccaca acgaggacta caccatcgtg gaacagtacg

2161 aacgcgccga gggccgccac tccaccggcg gcatggacga gctgtacaag taagaattcg

2221 atatcaagct tatcgataat caacctctgg attacaaaat ttgtgaaaga ttgactggta

2281 ttcttaacta tgttgctcct tttacgctat gtggatacgc tgctttaatg cctttgtatc

2341 atgctattgc ttcccgtatg gctttcattt tctcctcctt gtataaatcc tggttgctgt

2401 ctctttatga ggagttgtgg cccgttgtca ggcaacgtgg cgtggtgtgc actgtgtttg

2461 ctgacgcaac ccccactggt tggggcattg ccaccacctg tcagctcctt tccggaactt

2521 tcgctttccc cctccctatt gccacggcgg aactcatcgc cgcctgcctt gcccgctgct

2581 ggacaggggc tcggctgttg ggcactgaca attccgtggt gttgtcgggg aaatcatcgt

2641 cctttccttg gctgctcgcc tatgttgcca cctggattct gcgcgggacg tccttctgct

2701 acgtcccttc ggccctcaat ccagcggacc ttccttcccg cggcctgctg ccggctctgc

2761 ggcctcttcc gcgtcttcgc cttcgccctc agacgagtcg gatctccctt tgggccgcct

2821 ccccgcatcg ataccgagcg ctgctcgaga gatctacggg tggcatccct gtgacccctc

2881 cccagtgcct ctcctggccc tggaagttgc cactccagtg cccaccagcc ttgtcctaat

2941 aaaattaagt tgcatcattt tgtctgacta ggtgtccttc tataatatta tggggtggag

3001 gggggtggta tggagcaagg ggcaagttgg gaagacaacc tgtagggcct gcggggtcta

3061 ttgggaacca agctggagtg cagtggcaca atcttggctc actgcaatct ccgcctcctg

3121 ggttcaagcg attctcctgc ctcagcctcc cgagttgttg ggattccagg catgcatgac

3181 caggctcagc taatttttgt ttttttggta gagacggggt ttcaccatat tggccaggct

3241 ggtctccaac tcctaatctc aggtgatcta cccaccttgg cctcccaaat tgctgggatt

3301 acaggcgtga accactgctc ccttccctgt ccttctgatt ttgtaggtaa ccacgtgcgg

3361 accgagcggc cgcaggaacc cctagtgatg gagttggcca ctccctctct gcgcgctcgc

3421 tcgctcactg aggccgggcg accaaaggtc gcccgacgcc ggggctttgc ccgggcggcc

3481 tcagtgagcg agcgagcgcg cagagaggga gtggccaaa

//

### AAV9-CaMKIIα-hM4Di-mCherry

#### Summary Report

1-mer (%) 2-mer (%)

moles	100.0	0.0

mass	100.0	0.0

*************************

#### Gel Image

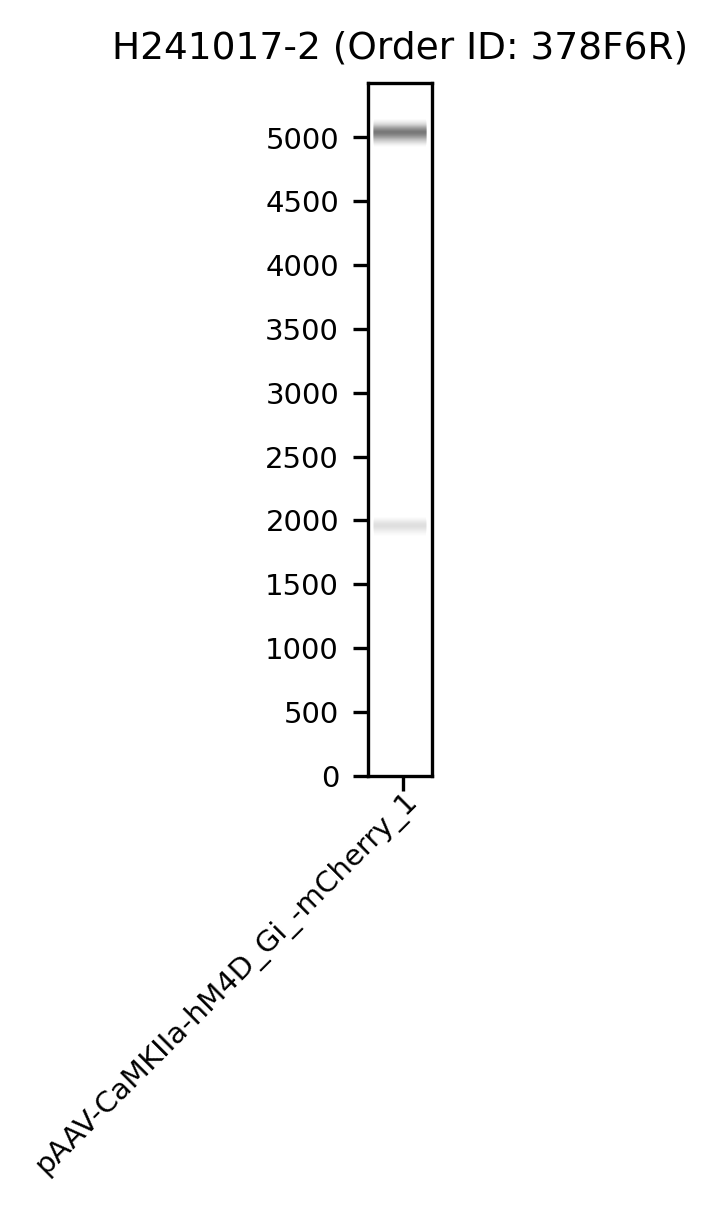

#### Histogram

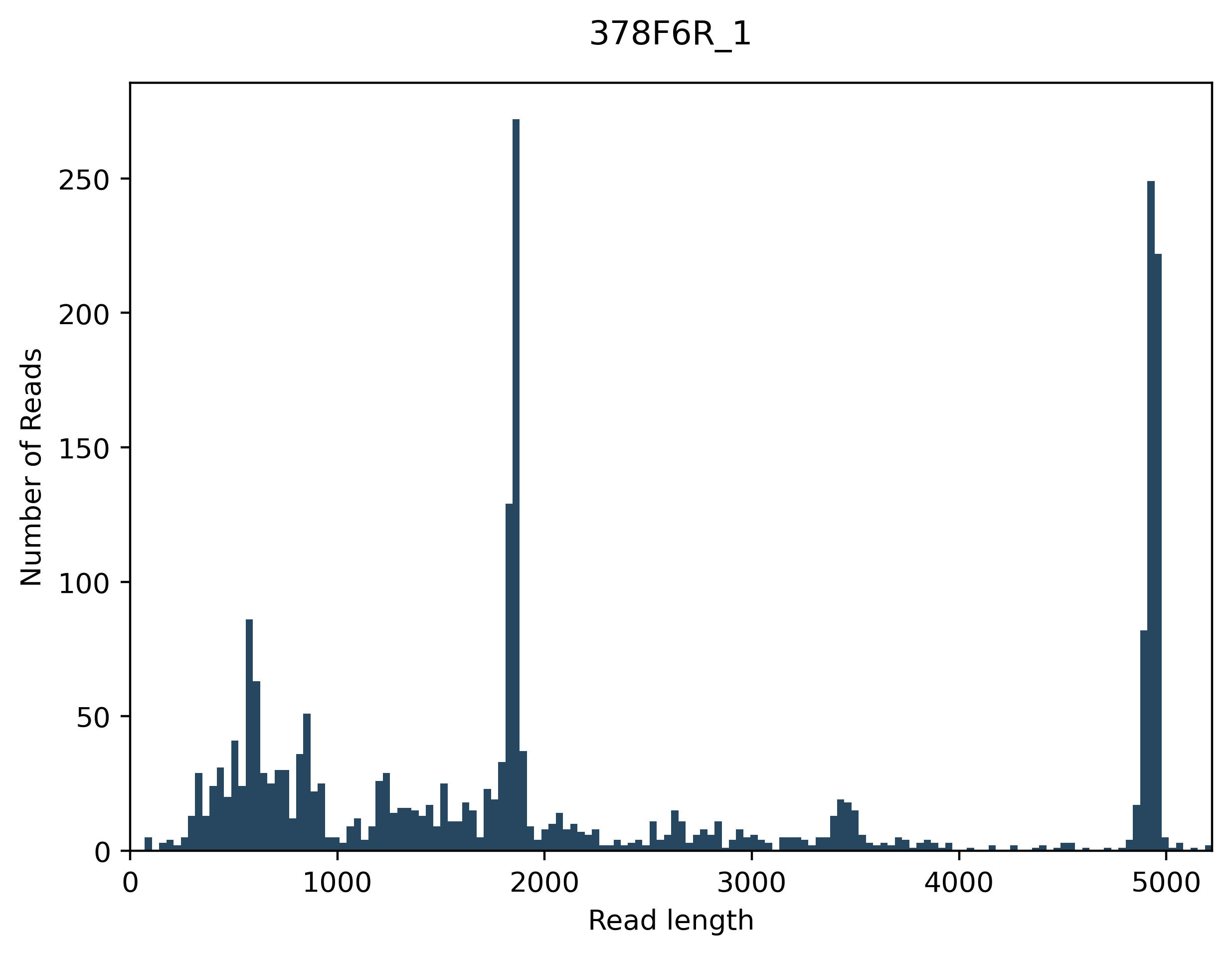

#### FASTA Assemblies

File: 378F6R_1_assembly-1_pAAV-CaMKIIa-hM4D_Gi_-mCherry.fasta

>pAAV-CaMKIIa-hM4D_Gi_-mCherry

CCACTCCCTCTCTGCGCGCTCGCTCGCTCACTGAGGCCGCCCGAGCAAAGCCCGCGCGTC

GGGCGACCTTTGCTCGCCCGGCCTCAGTGAGCGAGCGAGCGCGCAGAGAGGGAGTGGCCA

ACTCCATCACTAGGGGTTCCTGCGGCCGCACGCGTTTAACATTATGGCCTTAGGTCACTT

CATCTCCATGGGGTTCTTCTTCTGATTTTCTAGAAAATGAGATGGGGGTGCAGAGAGCTT

CCTCAGTGACCTGCCCAGGGTCACATCAGAAATGTCAGAGCTAGAACTTGAACTCAGATT

ACTAATCTTAAATTCCATGCCTTGGGGGCATGCAAGTACGATATACAGAAGGAGTGAACT

CATTAGGGCAGATGACCAATGAGTTTAGGAAAGAAGAGTCCAGGGCAGGGTACATCTACA

CCACCCGCCCAGCCCTGGGTGAGTCCAGCCACGTTCACCTCATTATAGTTGCCTCTCTCC

AGTCCTACCTTGACGGGAAGCACAAGCAGAAACTGGGACAGGAGCCCCAGGAGACCAAAT

CTTCATGGTCCCTCTGGGAGGATGGGTGGGGAGAGCTGTGGCAGAGGCCTCAGGAGGGGC

CCTGCTGCTCAGTGGTGACAGATAGGGGTGAGAAAGCAGACAGAGTCATTCCGTCAGCAT

TCTGGGTCTGTTTGGTACTTCTTCTCACGCTAAGGTGGCGGTGTGATATGCACAATGGCT

AAAAAGCAGGGAGAGCTGGAAAGAAACAAGGACAGAGACAGAGGCCAAGTCAACCAGACC

AATTCCCAGAGGAAGCAAAGAAACCATTACAGAGACTACAAGGGGGAAGGGAAGGAGAGA

TGAATTAGCTTCCCCTGTAAACCTTAGAACCCAGCTGTTGCCAGGGCAACGGGGCAATAC

CTGTCTCTTCAGAGGAGATGAAGTTGCCAGGGTAACTACATCCTGTCTTTCTCAAGGACC

ATCCCAGAATGTGGCACCCACTAGCCGTTACCATAGCAACTGCCTCTTTGCCCCACTTAA

TCCCATCCCGTCTGTTAAAAGGGCCCTATAGTTGGAGGTGGGGGAGGTAGGAAGAGCGAT

GATCACTTGTGGACTAAGTTTGTTCGCATCCCCTTCTCCAACCCCCTCAGTACATCACCC

TGGGGGAACAGGGTCCACTTGCTCCTGGGCCCACACAGTCCTGCAGTATTGTGTATATAA

GGCCAGGGCAAAGAGGAGCAGGTTTTAAAGTGAAAGGCAGGCAGGTGTTGGGGAGGCAGT

TACCGGGGCAACGGGAACAGGGCGTTTCGGAGGTGGTTGCCATGGGGACCTGGATGCTGA

CGAAGGCTCGCGAGGCTGTGAGCAGCCACAGTGCCCTGCTCAGAAGCCCCAAGCTCGTCA

GTCAAGCCGGTTCTCCGTTTGCACTCAGGAGCACGGGCAGGCGAGTGGCCCCTAGTTCTG

GGGGCAGCTCTAGAGCGGTACCGGATCCTCTAGAGTCGACGCCACCATGGCCAACTTCAC

ACCTGTCAATGGCAGCTCGGGCAATCAGTCCGTGCGCCTGGTCACGTCATCATCCCACAA

TCGCTATGAGACGGTGGAAATGGTCTTCATTGCCACAGTGACAGGCTCCCTGAGCCTGGT

GACTGTCGTGGGCAACATCCTGGTGATGCTGTCCATCAAGGTCAACAGGCAGCTGCAGAC

AGTCAACAACTACTTCCTCTTCAGCCTGGCGTGTGCTGATCTCATCATAGGCGCCTTCTC

CATGAACCTCTACACCGTGTACATCATCAAGGGCTACTGGCCCCTGGGCGCCGTGGTCTG

CGACCTGTGGCTGGCCCTGGACTGCGTGGTGAGCAACGCCTCCGTCATGAACCTTCTCAT

CATCAGCTTTGACCGCTACTTCTGCGTCACCAAGCCTCTCACCTACCCTGCCCGGCGCAC

CACCAAGATGGCAGGCCTCATGATTGCTGCTGCCTGGGTACTGTCCTTCGTGCTCTGGGC

GCCTGCCATCTTGTTCTGGCAGTTTGTGGTGGGTAAGCGGACGGTGCCCGACAACCAGTG

CTTCATCCAGTTCCTGTCCAACCCAGCAGTGACCTTTGGCACAGCCATTGCTGGCTTCTA

CCTGCCTGTGGTCATCATGACGGTGCTGTACATCCACATCTCCCTGGCCAGTCGCAGCCG

AGTCCACAAGCACCGGCCCGAGGGCCCGAAGGAGAAGAAAGCCAAGACGCTGGCCTTCCT

CAAGAGCCCACTAATGAAGCAGAGCGTCAAGAAGCCCCCGCCCGGGGAGGCCGCCCGGGA

GGAGCTGCGCAATGGCAAGCTGGAGGAGGCCCCCCCGCCAGCGCTGCCACCGCCACCGCG

CCCCGTGGCTGATAAGGACACTTCCAATGAGTCCAGCTCAGGCAGTGCCACCCAGAACAC

CAAGGAACGCCCAGCCACAGAGCTGTCCACCACAGAGGCCACCACGCCCGCCATGCCCGC

CCCTCCCCTGCAGCCGCGGGCCCTCAACCCAGCCTCCAGATGGTCCAAGATCCAGATTGT

GACGAAGCAGACAGGCAATGAGTGTGTGACAGCCATTGAGATTGTGCCTGCCACGCCGGC

TGGCATGCGCCCTGCGGCCAACGTGGCCCGCAAGTTCGCCAGCATCGCTCGCAACCAGGT

GCGCAAGAAGCGGCAGATGGCGGCCCGGGAGCGCAAAGTGACACGAACGATCTTTGCCAT

TCTGCTGGCCTTCATCCTCACCTGGACGCCCTACAACGTCATGGTCCTGGTGAACACCTT

CTGCCAGAGCTGCATCCCTGACACGGTGTGGTCCATTGGCTACTGGCTCTGCTACGTCAA

CAGCACCATCAACCCTGCCTGCTATGCTCTGTGCAACGCCACCTTTAAAAAGACCTTCCG

GCACCTGCTGCTGTGCCAGTATCGGAACATCGGCACTGCCAGGCGGGATCCACCGGTCGC

CACCATGGTGAGCAAGGGCGAGGAGGATAACATGGCCATCATCAAGGAGTTCATGCGCTT

CAAGGTGCACATGGAGGGCTCCGTGAACGGCCACGAGTTCGAGATCGAGGGCGAGGGCGA

GGGCCGCCCCTACGAGGGCACCCAGACCGCCAAGCTGAAGGTGACCAAGGGTGGCCCCCT

GCCCTTCGCCTGGGACATCCTGTCCCCTCAGTTCATGTACGGCTCCAAGGCCTACGTGAA

GCACCCCGCCGACATCCCCGACTACTTGAAGCTGTCCTTCCCCGAGGGCTTCAAGTGGGA

GCGCGTGATGAACTTCGAGGACGGCGGCGTGGTGACCGTGACCCAGGACTCCTCCCTGCA

GGACGGCGAGTTCATCTACAAGGTGAAGCTGCGCGGCACCAACTTCCCCTCCGACGGCCC

CGTAATGCAGAAGAAGACCATGGGCTGGGAGGCCTCCTCCGAGCGGATGTACCCCGAGGA

CGGCGCCCTGAAGGGCGAGATCAAGCAGAGGCTGAAGCTGAAGGACGGCGGCCACTACGA

CGCTGAGGTCAAGACCACCTACAAGGCCAAGAAGCCCGTGCAGCTGCCCGGCGCCTACAA

CGTCAACATCAAGTTGGACATCACCTCCCACAACGAGGACTACACCATCGTGGAACAGTA

CGAACGCGCCGAGGGCCGCCACTCCACCGGCGGCATGGACGAGCTGTACAAGTAAGAATT

CGATATCAAGCTTATCGATAATCAACCTCTGGATTACAAAATTTGTGAAAGATTGACTGG

TATTCTTAACTATGTTGCTCCTTTTACGCTATGTGGATACGCTGCTTTAATGCCTTTGTA

TCATGCTATTGCTTCCCGTATGGCTTTCATTTTCTCCTCCTTGTATAAATCCTGGTTGCT

GTCTCTTTATGAGGAGTTGTGGCCCGTTGTCAGGCAACGTGGCGTGGTGTGCACTGTGTT

TGCTGACGCAACCCCCACTGGTTGGGGCATTGCCACCACCTGTCAGCTCCTTTCCGGAAC

TTTCGCTTTCCCCCTCCCTATTGCCACGGCGGAACTCATCGCCGCCTGCCTTGCCCGCTG

CTGGACAGGGGCTCGGCTGTTGGGCACTGACAATTCCGTGGTGTTGTCGGGGAAATCATC

GTCCTTTCCTTGGCTGCTCGCCTATGTTGCCACCTGGATTCTGCGCGGGACGTCCTTCTG

CTACGTCCCTTCGGCCCTCAATCCAGCGGACCTTCCTTCCCGCGGCCTGCTGCCGGCTCT

GCGGCCTCTTCCGCGTCTTCGCCTTCGCCCTCAGACGAGTCGGATCTCCCTTTGGGCCGC

CTCCCCGCATCGATACCGAGCGCTGCTCGAGAGATCTACGGGTGGCATCCCTGTGACCCC

TCCCCAGTGCCTCTCCTGGCCCTGGAAGTTGCCACTCCAGTGCCCACCAGCCTTGTCCTA

ATAAAATTAAGTTGCATCATTTTGTCTGACTAGGTGTCCTTCTATAATATTATGGGGTGG

AGGGGGGTGGTATGGAGCAAGGGGCAAGTTGGGAAGACAACCTGTAGGGCCTGCGGGGTC

TATTGGGAACCAAGCTGGAGTGCAGTGGCACAATCTTGGCTCACTGCAATCTCCGCCTCC

TGGGTTCAAGCGATTCTCCTGCCTCAGCCTCCCGAGTTGTTGGGATTCCAGGCATGCATG

ACCAGGCTCAGCTAATTTTTGTTTTTTTGGTAGAGACGGGGTTTCACCATATTGGCCAGG

CTGGTCTCCAACTCCTAATCTCAGGTGATCTACCCACCTTGGCCTCCCAAATTGCTGGGA

TTACAGGCGTGAACCACTGCTCCCTTCCCTGTCCTTCTGATTTTGTAGGTAACCACGTGC

GGACCGAGCGGCCGCAGGAACCCCTAGTGATGGAGTTGGCCACTCCCTCTCTGCGCGCTC

GCTCGCTCACTGAGGCCGCGCGGCCAAAGGTCGCCGGGCGTCGGGGACCTTTGCTCGCCC

GGCCTCAGTGAGCGAGCGAGCGCGCAGA

File: 378F6R_1_assembly-2_pAAV-CaMKIIa-hM4D_Gi_-mCherry.fasta

>pAAV-CaMKIIa-hM4D_Gi_-mCherry

TTGGCCACTCCCTCTCTGCGCGCTCGCTCGCTCACTGAGGCCGGGCGACCAAAGGTCGCC

CGACGCCCGGGCTTTGCCCGGGCGGCCTCAGTGAGCGAGCGAGCGCGCAGAGAGGGAGTG

GCCAACTCCATCACTAGGGGTTCCTGCGGCCGCTCGGTCCGCACGTGGTTACCTACAAAA

TCAGAAGGACAGGGAAGGGAGCAGTGGTTCACGCCTGTAATCCCAGCAATTTGGGAGGCC

AAGGTGGGTAGATCACCTGAGATTAGGAGTTGGAGACCAGCCTGGCCAATATGGTGAAAC

CCCGTCTCTACCAAAAAAACAAAAATTAGCTGAGCCTGGTCATGCATGCCTGGAATCCCA

ACAACTCGGGAGGCTGAGGCAGGAGAATCGCTTGAACCCAGGAGGCGGAGATTGCAGTGA

GCCAAGATTGTGCCACTGCACTCCAGCTTGGTTCCCAATAGACCCCGCAGGCCCTACAGG

TTGTCTTCCCAACTTGCCCCTTGCTCCATACCACCCCCCTCCACCCCATAATATTATAGA

AGGACACCTAGTCAGACAAAATGATGCAACTTAATTTTATTAGGACAAGGCTGGTGGGCA

CTGGAGTGGCAACTTCCAGGGCCAGGAGAGGCACTGGGGAGGGGTCACAGGGATGCCACC

CGTAGATCTCTCGAGCAGCGCTCGGTATCGATGCGGGGAGGCGGCCCAAAGGGAGATCCG

ACTCGTCTGAGGGCGAAGGCGAAGACGCGGAAGAGGCCGCAGAGCCGGCAGCAGGCCGCG

GGAAGGAAGGTCCGCTGGATTGAGGGCCGAAGGGACGTAGCAGAAGGACGTCCCGCGCAG

AATCCAGGTGGCAACATAGGCGAGCAGCCAAGGAAAGGACGATGATTTCCCCGACAACAC

CACGGAATTGTCAGTGCCCAACAGCCGAGCCCCTGTCCAGCAGCGGGCAAGGCAGGCGGC

GATGAGTTCCGCCGTGGTGTTGTCGGGGAAATCATCGTCCTTTCCTTGGCTGCTCGCCTA

TGTTGCCACCTGGATTCTGCGCGGGACGTCCTTCTGCTACGTCCCTTCGGCCCTCAATCC

AGCGGACCTTCCTTCCCGCGGCCTGCTGCCGGCTCTGCGGCCTCTTCCGCGTCTTCGCCT

TCGCCCTCAGACGAGTCGGATCTCCCTTTGGGCCGCCTCCCCGCATCGATACCGAGCGCT

GCTCGAGAGATCTACGGGTGGCATCCCTGTGACCCCTCCCCAGTGCCTCTCCTGGCCCTG

GAAGTTGCCACTCCAGTGCCCACCAGCCTTGTCCTAATAAAATTAAGTTGCATCATTTTG

TCTGACTAGGTGTCCTTCTATAATATTATGGGGTGGAGGGGGGTGGTATGGAGCAAGGGG

CAAGTTGGGAAGACAACCTGTAGGGCCTGCGGGGTCTATTGGGAACCAAGCTGGAGTGCA

GTGGCACAATCTTGGCTCACTGCAATCTCCGCCTCCTGGGTTCAAGCGATTCTCCTGCCT

CAGCCTCCCGAGTTGTTGGGATTCCAGGCATGCATGACCAGGCTCAGCTAATTTTTGTTT

TTTTGGTAGAGACGGGGTTTCACCATATTGGCCAGGCTGGTCTCCAACTCCTAATCTCAG

GTGATCTACCCACCTTGGCCTCCCAAATTGCTGGGATTACAGGCGTGAACCACTGCTCCC

TTCCCTGTCCTTCTGATTTTGTAGGTAACCACGTGCGGACCGAGCGGCCGCAGGAACCCC

TAGTGATGGAGTTGGCCACTCCCTCTCTGCGCGCTCGCTCGCTCACTGAGGCCGGGCGAC

CAAAGGTCGCCCGACGCCGGGGCTTTGCCCGGGCGGCCTCAGTGAGCGAGCGAGCGCGCA

GAGAGGGAGTGGCCAAA

File: 378F6R_1_assembly-3_pAAV-CaMKIIa-hM4D_Gi_-mCherry.fasta

>pAAV-CaMKIIa-hM4D_Gi_-mCherry

TTGGCCACTCCCTCTCTGCGCGCTCGCTCGCTCACTGAGGCCGCCCGGGCAAAGCCCGGG

CGTCGGGCGACCTTTGCTCGCCCGGCCTCAGTGAGCGAGCGAGCGCGCAGAGAGGGAGTG

GCCAACTCCATCACTAGGGGTTCCTGCGGCCGCTCGGTCCGCACGTGGTTACCTACAAAA

TCAGAAGGACAGGGAAGGGAGCAGTGGTTCACGCCTGTAATCCCAGCAATTTGGGAGGCC

AAGGTGGGTAGATCACCTGAGATTAGGAGTTGGAGACCAGCCTGGCCAATATGGTGAAAC

CCCGTCTCTACCAAAAAAACAAAAATTAGCTGAGCCTGGTCATGCATGCCTGGAATCCCA

ACAACTCGGGAGGCTGAGGCAGGAGAATCGCTTGAACCCAGGAGGCGGAGATTGCAGTGA

GCCAAGATTGTGCCACTGCACTCCAGCTTGGTTCCCAATAGACCCCGCAGGCCCTACAGG

TTGTCTTCCCAACTTGCCCCTTGCTCCATACCACCCCCCTCCACCCCATAATATTATAGA

AGGACACCTAGTCAGACAAAATGATGCAACTTAATTTTATTAGGACAAGGCTGGTGGGCA

CTGGAGTGGCAACTTCCAGGGCCAGGAGAGGCACTGGGGAGGGGTCACAGGGATGCCACC

CGTAGATCTCTCGAGCAGCGCTCGGTATCGATGCGGGGAGGCGGCCCAAAGGGAGATCCG

ACTCGTCTGAGGGCGAAGGCGAAGACGCGGAAGAGGCCGCAGAGCCGGCAGCAGGCCGCG

GGAAGGAAGGTCCGCTGGATTGAGGGCCGAAGGGACGTAGCAGAAGGACGTCCCGCGCAG

AATCCAGGTGGCAACATAGGCGAGCAGCCAAGGAAAGGACGATGATTTCCCCGACAACAC

CACGGAATTGTCAGTGCCCAACAGCCGAGCCCCTGTCCAGCAGCGGGCAAGGCAGGCGGC

GATGAGTTCCGCCGTGGCAATAGGGAGGGGGAAAGCGAAAGTTCCGGAAAGGAGCTGACA

GGTGGTGGCAATGCCCCAACCAGTGGGGGTTGCGTCAGCAAACACAGTGCACACCACGCC

ACGTTGCCTGACAACGGGCCACAACTCCTCATAAAGAGACAGCAACCAGGATTTATACAA

GGAGGAGAAAATGAAAGCCATACGGGAAGCAATAGCATGATACAAAGGCATTAAAGCAGC

GTATCCACATAGCGTAAAAGGAGCAACATAGTTAAGAATACCAGTCAATCTTTCACAAAT

TTTGTAATCCAGAGGTTGATTATCGATAAGCTTGATATCGAATTCTTACTTGTACAGCTC

GTCCATGCCGCCGGTGGAGTGGCGGCCCTCGGCGCGTTCGTACTGTTCCACGATGGTGTA

GTCCTCGTTGTGGGAGGTGATGTCCAACTTGATGTTGACGTTGTAGGCGCCGGGCAGCTG

CACGGGCTTCTTGGCCTTGTAGGTGGTCTTGACCTCAGCGTCGTAGTGGCCGCCGTCCTT

CAGCTTCAGCCTCTGCTTGATCTCGCCCTTCAGGGCGCCGTCCTCGGGGTACATCCGCTC

GGAGGAGGCCTCCCAGCCCATGGTCTTCTTCTGCATTACGGGGCCGTCGGAGGGGAAGTT

GGTGCCGCGCAGCTTCACCTTGTAGATGAACTCGCCGTCCTGCAGGGAGGAGTCCTGGGT

CACGGTCACCACGCCGCCGTCCCCTCAGTTCATTACGGCTCCAAGGCCTACGTGAAGCAC

CCCGCCGACATCCCCGACTACTTGAAGCTGTCCTTCCCCGAGGGCTTCAAGTGGGAGCGC

GTGATGAACTTCGAGGACGGCGGCGTGGTGACCGTGACCCAGGACTCCTCCCTGCAGGAC

GGCGAGTTCATCTACAAGGTGAAGCTGCGCGGCACCAACTTCCCCTCCGACGGCCCCGTA

ATGCAGAAGAAGACCATGGGCTGGGAGGCCTCCTCCGAGCGGATGTACCCCGAGGACGGC

GCCCTGAAGGGCGAGATCAAGCAGAGGCTGAAGCTGAAGGACGGCGGCCACTACGACGCT

GAGGTCAAGACCACCTACAAGGCCAAGAAGCCCGTGCAGCTGCCCGGCGCCTACAACGTC

AACATCAAGTTGGACATCACCTCCCACAACGAGGACTACACCATCGTGGAACAGTACGAA

CGCGCCGAGGGCCGCCACTCCACCGGCGGCATGGACGAGCTGTACAAGTAAGAATTCGAT

ATCAAGCTTATCGATAATCAACCTCTGGATTACAAAATTTGTGAAAGATTGACTGGTATT

CTTAACTATGTTGCTCCTTTTACGCTATGTGGATACGCTGCTTTAATGCCTTTGTATCAT

GCTATTGCTTCCCGTATGGCTTTCATTTTCTCCTCCTTGTATAAATCCTGGTTGCTGTCT

CTTTATGAGGAGTTGTGGCCCGTTGTCAGGCAACGTGGCGTGGTGTGCACTGTGTTTGCT

GACGCAACCCCCACTGGTTGGGGCATTGCCACCACCTGTCAGCTCCTTTCCGGAACTTTC

GCTTTCCCCCTCCCTATTGCCACGGCGGAACTCATCGCCGCCTGCCTTGCCCGCTGCTGG

ACAGGGGCTCGGCTGTTGGGCACTGACAATTCCGTGGTGTTGTCGGGGAAATCATCGTCC

TTTCCTTGGCTGCTCGCCTATGTTGCCACCTGGATTCTGCGCGGGACGTCCTTCTGCTAC

GTCCCTTCGGCCCTCAATCCAGCGGACCTTCCTTCCCGCGGCCTGCTGCCGGCTCTGCGG

CCTCTTCCGCGTCTTCGCCTTCGCCCTCAGACGAGTCGGATCTCCCTTTGGGCCGCCTCC

CCGCATCGATACCGAGCGCTGCTCGAGAGATCTACGGGTGGCATCCCTGTGACCCCTCCC

CAGTGCCTCTCCTGGCCCTGGAAGTTGCCACTCCAGTGCCCACCAGCCTTGTCCTAATAA

AATTAAGTTGCATCATTTTGTCTGACTAGGTGTCCTTCTATAATATTATGGGGTGGAGGG

GGGTGGTATGGAGCAAGGGGCAAGTTGGGAAGACAACCTGTAGGGCCTGCGGGGTCTATT

GGGAACCAAGCTGGAGTGCAGTGGCACAATCTTGGCTCACTGCAATCTCCGCCTCCTGGG

TTCAAGCGATTCTCCTGCCTCAGCCTCCCGAGTTGTTGGGATTCCAGGCATGCATGACCA

GGCTCAGCTAATTTTTGTTTTTTTGGTAGAGACGGGGTTTCACCATATTGGCCAGGCTGG

TCTCCAACTCCTAATCTCAGGTGATCTACCCACCTTGGCCTCCCAAATTGCTGGGATTAC

AGGCGTGAACCACTGCTCCCTTCCCTGTCCTTCTGATTTTGTAGGTAACCACGTGCGGAC

CGAGCGGCCGCAGGAACCCCTAGTGATGGAGTTGGCCACTCCCTCTCTGCGCGCTCGCTC

GCTCACTGAGGCCGCGCGGGCAAAGGCCCCGGCGCCGGGCGACCTTTGCTCGCGCGGCCT

CAGTGAGCGAGCGAGCGCGCAGAGAGGGAGTGGCCAAA

#### GenBank Annotations

File: 378F6R_1_assembly-1_pAAV-CaMKIIa-hM4D_Gi_-mCherry.gbk

LOCUS	pAAV-CaMKIIa-hM4D_Gi_-mCherry 4948 bp	DNA	linear	SYN 07-NOV-2024

DEFINITION pAAV-CaMKIIa-hM4D_Gi_-mCherry.

ACCESSION	pAAV-CaMKIIa-hM4D_Gi_-mCherry

VERSION	pAAV-CaMKIIa-hM4D_Gi_-mCherry

KEYWORDS	.

SOURCE

ORGANISM .

.

COMMENT	Annotated with pLannotate v1.2.2

FEATURES	Location/Qualifiers

promoter	160..1448

/note="pLannotate"

/label="CaMKII promoter"

/database="snapgene"

/identity="100.0"

/match_length="100.0"

/fragment="False"

/other="promoter"

CDS	2945..3652

/note="pLannotate"

/label="mCherry"

/database="fpbase"

/identity="100.0"

/match_length="100.0"

/fragment="False"

/other="CDS"

polyA_signal	4300..4776

/note="pLannotate"

/label="hGH poly(A) signal"

/database="snapgene"

/identity="100.0"

/match_length="100.0"

/fragment="False"

/other="polyA_signal"

CDS	1487..2923

/note="pLannotate"

/label="CHRM4"

/database="swissprot"

/identity="99.6"

/match_length="100.0"

/fragment="False"

/other="CDS"

misc_feature	3680..4268

/note="pLannotate"

/label="WPRE"

/database="snapgene"

/identity="99.7"

/match_length="100.0"

/fragment="False"

/other="misc_feature"

repeat_region	complement(12..141)

/note="pLannotate"

/label="AAV2 ITR (fragment)"

/database="snapgene"

/identity="97.7"

/match_length="92.2"

/fragment="True"

/other="repeat_region"

misc_feature	3656..3680

/note="pLannotate"

/label="MCS (fragment)"

/database="snapgene"

/identity="100.0"

/match_length="23.1"

/fragment="True"

/other="misc_feature"

misc_feature	1463..1480

/note="pLannotate"

/label="MCS (fragment)"

/database="snapgene"

/identity="100.0"

/match_length="31.6"

/fragment="True"

/other="misc_feature"

ORIGIN

1 ccactccctc tctgcgcgct cgctcgctca ctgaggccgc ccgagcaaag cccgcgcgtc

61 gggcgacctt tgctcgcccg gcctcagtga gcgagcgagc gcgcagagag ggagtggcca

121 actccatcac taggggttcc tgcggccgca cgcgtttaac attatggcct taggtcactt

181 catctccatg gggttcttct tctgattttc tagaaaatga gatgggggtg cagagagctt

241 cctcagtgac ctgcccaggg tcacatcaga aatgtcagag ctagaacttg aactcagatt

301 actaatctta aattccatgc cttgggggca tgcaagtacg atatacagaa ggagtgaact

361 cattagggca gatgaccaat gagtttagga aagaagagtc cagggcaggg tacatctaca

421 ccacccgccc agccctgggt gagtccagcc acgttcacct cattatagtt gcctctctcc

481 agtcctacct tgacgggaag cacaagcaga aactgggaca ggagccccag gagaccaaat

541 cttcatggtc cctctgggag gatgggtggg gagagctgtg gcagaggcct caggaggggc

601 cctgctgctc agtggtgaca gataggggtg agaaagcaga cagagtcatt ccgtcagcat

661 tctgggtctg tttggtactt cttctcacgc taaggtggcg gtgtgatatg cacaatggct

721 aaaaagcagg gagagctgga aagaaacaag gacagagaca gaggccaagt caaccagacc

781 aattcccaga ggaagcaaag aaaccattac agagactaca agggggaagg gaaggagaga

841 tgaattagct tcccctgtaa accttagaac ccagctgttg ccagggcaac ggggcaatac

901 ctgtctcttc agaggagatg aagttgccag ggtaactaca tcctgtcttt ctcaaggacc

961 atcccagaat gtggcaccca ctagccgtta ccatagcaac tgcctctttg ccccacttaa

1021 tcccatcccg tctgttaaaa gggccctata gttggaggtg ggggaggtag gaagagcgat

1081 gatcacttgt ggactaagtt tgttcgcatc cccttctcca accccctcag tacatcaccc

1141 tgggggaaca gggtccactt gctcctgggc ccacacagtc ctgcagtatt gtgtatataa

1201 ggccagggca aagaggagca ggttttaaag tgaaaggcag gcaggtgttg gggaggcagt

1261 taccggggca acgggaacag ggcgtttcgg aggtggttgc catggggacc tggatgctga

1321 cgaaggctcg cgaggctgtg agcagccaca gtgccctgct cagaagcccc aagctcgtca

1381 gtcaagccgg ttctccgttt gcactcagga gcacgggcag gcgagtggcc cctagttctg

1441 ggggcagctc tagagcggta ccggatcctc tagagtcgac gccaccatgg ccaacttcac

1501 acctgtcaat ggcagctcgg gcaatcagtc cgtgcgcctg gtcacgtcat catcccacaa

1561 tcgctatgag acggtggaaa tggtcttcat tgccacagtg acaggctccc tgagcctggt

1621 gactgtcgtg ggcaacatcc tggtgatgct gtccatcaag gtcaacaggc agctgcagac

1681 agtcaacaac tacttcctct tcagcctggc gtgtgctgat ctcatcatag gcgccttctc

1741 catgaacctc tacaccgtgt acatcatcaa gggctactgg cccctgggcg ccgtggtctg

1801 cgacctgtgg ctggccctgg actgcgtggt gagcaacgcc tccgtcatga accttctcat

1861 catcagcttt gaccgctact tctgcgtcac caagcctctc acctaccctg cccggcgcac

1921 caccaagatg gcaggcctca tgattgctgc tgcctgggta ctgtccttcg tgctctgggc

1981 gcctgccatc ttgttctggc agtttgtggt gggtaagcgg acggtgcccg acaaccagtg

2041 cttcatccag ttcctgtcca acccagcagt gacctttggc acagccattg ctggcttcta

2101 cctgcctgtg gtcatcatga cggtgctgta catccacatc tccctggcca gtcgcagccg

2161 agtccacaag caccggcccg agggcccgaa ggagaagaaa gccaagacgc tggccttcct

2221 caagagccca ctaatgaagc agagcgtcaa gaagcccccg cccggggagg ccgcccggga

2281 ggagctgcgc aatggcaagc tggaggaggc ccccccgcca gcgctgccac cgccaccgcg

2341 ccccgtggct gataaggaca cttccaatga gtccagctca ggcagtgcca cccagaacac

2401 caaggaacgc ccagccacag agctgtccac cacagaggcc accacgcccg ccatgcccgc

2461 ccctcccctg cagccgcggg ccctcaaccc agcctccaga tggtccaaga tccagattgt

2521 gacgaagcag acaggcaatg agtgtgtgac agccattgag attgtgcctg ccacgccggc

2581 tggcatgcgc cctgcggcca acgtggcccg caagttcgcc agcatcgctc gcaaccaggt

2641 gcgcaagaag cggcagatgg cggcccggga gcgcaaagtg acacgaacga tctttgccat

2701 tctgctggcc ttcatcctca cctggacgcc ctacaacgtc atggtcctgg tgaacacctt

2761 ctgccagagc tgcatccctg acacggtgtg gtccattggc tactggctct gctacgtcaa

2821 cagcaccatc aaccctgcct gctatgctct gtgcaacgcc acctttaaaa agaccttccg

2881 gcacctgctg ctgtgccagt atcggaacat cggcactgcc aggcgggatc caccggtcgc

2941 caccatggtg agcaagggcg aggaggataa catggccatc atcaaggagt tcatgcgctt

3001 caaggtgcac atggagggct ccgtgaacgg ccacgagttc gagatcgagg gcgagggcga

3061 gggccgcccc tacgagggca cccagaccgc caagctgaag gtgaccaagg gtggccccct

3121 gcccttcgcc tgggacatcc tgtcccctca gttcatgtac ggctccaagg cctacgtgaa

3181 gcaccccgcc gacatccccg actacttgaa gctgtccttc cccgagggct tcaagtggga

3241 gcgcgtgatg aacttcgagg acggcggcgt ggtgaccgtg acccaggact cctccctgca

3301 ggacggcgag ttcatctaca aggtgaagct gcgcggcacc aacttcccct ccgacggccc

3361 cgtaatgcag aagaagacca tgggctggga ggcctcctcc gagcggatgt accccgagga

3421 cggcgccctg aagggcgaga tcaagcagag gctgaagctg aaggacggcg gccactacga

3481 cgctgaggtc aagaccacct acaaggccaa gaagcccgtg cagctgcccg gcgcctacaa

3541 cgtcaacatc aagttggaca tcacctccca caacgaggac tacaccatcg tggaacagta

3601 cgaacgcgcc gagggccgcc actccaccgg cggcatggac gagctgtaca agtaagaatt

3661 cgatatcaag cttatcgata atcaacctct ggattacaaa atttgtgaaa gattgactgg

3721 tattcttaac tatgttgctc cttttacgct atgtggatac gctgctttaa tgcctttgta

3781 tcatgctatt gcttcccgta tggctttcat tttctcctcc ttgtataaat cctggttgct

3841 gtctctttat gaggagttgt ggcccgttgt caggcaacgt ggcgtggtgt gcactgtgtt

3901 tgctgacgca acccccactg gttggggcat tgccaccacc tgtcagctcc tttccggaac

3961 tttcgctttc cccctcccta ttgccacggc ggaactcatc gccgcctgcc ttgcccgctg

4021 ctggacaggg gctcggctgt tgggcactga caattccgtg gtgttgtcgg ggaaatcatc

4081 gtcctttcct tggctgctcg cctatgttgc cacctggatt ctgcgcggga cgtccttctg

4141 ctacgtccct tcggccctca atccagcgga ccttccttcc cgcggcctgc tgccggctct

4201 gcggcctctt ccgcgtcttc gccttcgccc tcagacgagt cggatctccc tttgggccgc

4261 ctccccgcat cgataccgag cgctgctcga gagatctacg ggtggcatcc ctgtgacccc

4321 tccccagtgc ctctcctggc cctggaagtt gccactccag tgcccaccag ccttgtccta

4381 ataaaattaa gttgcatcat tttgtctgac taggtgtcct tctataatat tatggggtgg

4441 aggggggtgg tatggagcaa ggggcaagtt gggaagacaa cctgtagggc ctgcggggtc

4501 tattgggaac caagctggag tgcagtggca caatcttggc tcactgcaat ctccgcctcc

4561 tgggttcaag cgattctcct gcctcagcct cccgagttgt tgggattcca ggcatgcatg

4621 accaggctca gctaattttt gtttttttgg tagagacggg gtttcaccat attggccagg

4681 ctggtctcca actcctaatc tcaggtgatc tacccacctt ggcctcccaa attgctggga

4741 ttacaggcgt gaaccactgc tcccttccct gtccttctga ttttgtaggt aaccacgtgc

4801 ggaccgagcg gccgcaggaa cccctagtga tggagttggc cactccctct ctgcgcgctc

4861 gctcgctcac tgaggccgcg cggccaaagg tcgccgggcg tcggggacct ttgctcgccc

4921 ggcctcagtg agcgagcgag cgcgcaga

//

File: 378F6R_1_assembly-2_pAAV-CaMKIIa-hM4D_Gi_-mCherry.gbk

LOCUS	pAAV-CaMKIIa-hM4D_Gi_-mCherry 1877 bp	DNA	linear	SYN 07-NOV-2024

DEFINITION pAAV-CaMKIIa-hM4D_Gi_-mCherry.

ACCESSION	pAAV-CaMKIIa-hM4D_Gi_-mCherry

VERSION	pAAV-CaMKIIa-hM4D_Gi_-mCherry

KEYWORDS	.

SOURCE

ORGANISM .

.

COMMENT	Annotated with pLannotate v1.2.2

FEATURES	Location/Qualifiers

polyA_signal	1216..1692

/note="pLannotate"

/label="hGH poly(A) signal"

/database="snapgene"

/identity="100.0"

/match_length="100.0"

/fragment="False"

/other="polyA_signal"

polyA_signal	complement(185..661)

/note="pLannotate"

/label="hGH poly(A) signal"

/database="snapgene"

/identity="100.0"

/match_length="100.0"

/fragment="False"

/other="polyA_signal"

misc_feature	complement(693..977)

/note="pLannotate"

/label="WPRE (fragment)"

/database="snapgene"

/identity="99.6"

/match_length="48.4"

/fragment="True"

/other="misc_feature"

repeat_region	1732..1861

/note="pLannotate"

/label="AAV2 ITR (fragment)"

/database="snapgene"

/identity="99.2"

/match_length="92.2"

/fragment="True"

/other="repeat_region"

repeat_region	1..110

/note="pLannotate"

/label="AAV2 ITR (fragment)"

/database="snapgene"

/identity="100.0"

/match_length="78.0"

/fragment="True"

/other="repeat_region"

misc_feature	972..1184

/note="pLannotate"

/label="WPRE (fragment)"

/database="snapgene"

/identity="99.5"

/match_length="36.2"

/fragment="True"

/other="misc_feature"

ORIGIN

1 ttggccactc cctctctgcg cgctcgctcg ctcactgagg ccgggcgacc aaaggtcgcc

61 cgacgcccgg gctttgcccg ggcggcctca gtgagcgagc gagcgcgcag agagggagtg

121 gccaactcca tcactagggg ttcctgcggc cgctcggtcc gcacgtggtt acctacaaaa

181 tcagaaggac agggaaggga gcagtggttc acgcctgtaa tcccagcaat ttgggaggcc

241 aaggtgggta gatcacctga gattaggagt tggagaccag cctggccaat atggtgaaac

301 cccgtctcta ccaaaaaaac aaaaattagc tgagcctggt catgcatgcc tggaatccca

361 acaactcggg aggctgaggc aggagaatcg cttgaaccca ggaggcggag attgcagtga

421 gccaagattg tgccactgca ctccagcttg gttcccaata gaccccgcag gccctacagg

481 ttgtcttccc aacttgcccc ttgctccata ccacccccct ccaccccata atattataga

541 aggacaccta gtcagacaaa atgatgcaac ttaattttat taggacaagg ctggtgggca

601 ctggagtggc aacttccagg gccaggagag gcactgggga ggggtcacag ggatgccacc

661 cgtagatctc tcgagcagcg ctcggtatcg atgcggggag gcggcccaaa gggagatccg

721 actcgtctga gggcgaaggc gaagacgcgg aagaggccgc agagccggca gcaggccgcg

781 ggaaggaagg tccgctggat tgagggccga agggacgtag cagaaggacg tcccgcgcag

841 aatccaggtg gcaacatagg cgagcagcca aggaaaggac gatgatttcc ccgacaacac

901 cacggaattg tcagtgccca acagccgagc ccctgtccag cagcgggcaa ggcaggcggc

961 gatgagttcc gccgtggtgt tgtcggggaa atcatcgtcc tttccttggc tgctcgccta

1021 tgttgccacc tggattctgc gcgggacgtc cttctgctac gtcccttcgg ccctcaatcc

1081 agcggacctt ccttcccgcg gcctgctgcc ggctctgcgg cctcttccgc gtcttcgcct

1141 tcgccctcag acgagtcgga tctccctttg ggccgcctcc ccgcatcgat accgagcgct

1201 gctcgagaga tctacgggtg gcatccctgt gacccctccc cagtgcctct cctggccctg

1261 gaagttgcca ctccagtgcc caccagcctt gtcctaataa aattaagttg catcattttg

1321 tctgactagg tgtccttcta taatattatg gggtggaggg gggtggtatg gagcaagggg

1381 caagttggga agacaacctg tagggcctgc ggggtctatt gggaaccaag ctggagtgca

1441 gtggcacaat cttggctcac tgcaatctcc gcctcctggg ttcaagcgat tctcctgcct

1501 cagcctcccg agttgttggg attccaggca tgcatgacca ggctcagcta atttttgttt

1561 ttttggtaga gacggggttt caccatattg gccaggctgg tctccaactc ctaatctcag

1621 gtgatctacc caccttggcc tcccaaattg ctgggattac aggcgtgaac cactgctccc

1681 ttccctgtcc ttctgatttt gtaggtaacc acgtgcggac cgagcggccg caggaacccc

1741 tagtgatgga gttggccact ccctctctgc gcgctcgctc gctcactgag gccgggcgac

1801 caaaggtcgc ccgacgccgg ggctttgccc gggcggcctc agtgagcgag cgagcgcgca

1861 gagagggagt ggccaaa

//

File: 378F6R_1_assembly-3_pAAV-CaMKIIa-hM4D_Gi_-mCherry.gbk

LOCUS	pAAV-CaMKIIa-hM4D_Gi_-mCherry 3518 bp	DNA	linear	SYN 07-NOV-2024

DEFINITION pAAV-CaMKIIa-hM4D_Gi_-mCherry.

ACCESSION	pAAV-CaMKIIa-hM4D_Gi_-mCherry

VERSION	pAAV-CaMKIIa-hM4D_Gi_-mCherry

KEYWORDS	.

SOURCE

ORGANISM .

.

COMMENT	Annotated with pLannotate v1.2.2

FEATURES	Location/Qualifiers

polyA_signal	2856..3332

/note="pLannotate"

/label="hGH poly(A) signal"

/database="snapgene"

/identity="100.0"

/match_length="100.0"

/fragment="False"

/other="polyA_signal"

polyA_signal	complement(185..661)

/note="pLannotate"

/label="hGH poly(A) signal"

/database="snapgene"

/identity="100.0"

/match_length="100.0"

/fragment="False"

/other="polyA_signal"

misc_feature	2236..2824

/note="pLannotate"

/label="WPRE"

/database="snapgene"

/identity="99.7"

/match_length="100.0"

/fragment="False"

/other="misc_feature"

misc_feature	complement(693..1281)

/note="pLannotate"

/label="WPRE"

/database="snapgene"

/identity="99.7"

/match_length="100.0"

/fragment="False"

/other="misc_feature"

CDS	1651..2208

/note="pLannotate"

/label="Blue102 (fragment)"

/database="fpbase"

/identity="90.3"

/match_length="80.5"

/fragment="True"

/other="CDS"

CDS	complement(1306..1702)

/note="pLannotate"

/label="Slow-FT (fragment)"

/database="snapgene"

/identity="99.7"

/match_length="55.8"

/fragment="True"

/other="CDS"

repeat_region	complement(16..145)

/note="pLannotate"

/label="AAV2 ITR (fragment)"

/database="snapgene"

/identity="99.2"

/match_length="92.2"

/fragment="True"

/other="repeat_region"

misc_feature	2212..2236

/note="pLannotate"

/label="MCS (fragment)"

/database="snapgene"

/identity="100.0"

/match_length="23.1"

/fragment="True"

/other="misc_feature"

misc_feature	complement(1281..1305)

/note="pLannotate"

/label="MCS (fragment)"

/database="snapgene"

/identity="100.0"

/match_length="23.1"

/fragment="True"

/other="misc_feature"

ORIGIN

1 ttggccactc cctctctgcg cgctcgctcg ctcactgagg ccgcccgggc aaagcccggg

61 cgtcgggcga cctttgctcg cccggcctca gtgagcgagc gagcgcgcag agagggagtg

121 gccaactcca tcactagggg ttcctgcggc cgctcggtcc gcacgtggtt acctacaaaa

181 tcagaaggac agggaaggga gcagtggttc acgcctgtaa tcccagcaat ttgggaggcc

241 aaggtgggta gatcacctga gattaggagt tggagaccag cctggccaat atggtgaaac

301 cccgtctcta ccaaaaaaac aaaaattagc tgagcctggt catgcatgcc tggaatccca

361 acaactcggg aggctgaggc aggagaatcg cttgaaccca ggaggcggag attgcagtga

421 gccaagattg tgccactgca ctccagcttg gttcccaata gaccccgcag gccctacagg

481 ttgtcttccc aacttgcccc ttgctccata ccacccccct ccaccccata atattataga

541 aggacaccta gtcagacaaa atgatgcaac ttaattttat taggacaagg ctggtgggca

601 ctggagtggc aacttccagg gccaggagag gcactgggga ggggtcacag ggatgccacc

661 cgtagatctc tcgagcagcg ctcggtatcg atgcggggag gcggcccaaa gggagatccg

721 actcgtctga gggcgaaggc gaagacgcgg aagaggccgc agagccggca gcaggccgcg

781 ggaaggaagg tccgctggat tgagggccga agggacgtag cagaaggacg tcccgcgcag

841 aatccaggtg gcaacatagg cgagcagcca aggaaaggac gatgatttcc ccgacaacac

901 cacggaattg tcagtgccca acagccgagc ccctgtccag cagcgggcaa ggcaggcggc

961 gatgagttcc gccgtggcaa tagggagggg gaaagcgaaa gttccggaaa ggagctgaca

1021 ggtggtggca atgccccaac cagtgggggt tgcgtcagca aacacagtgc acaccacgcc

1081 acgttgcctg acaacgggcc acaactcctc ataaagagac agcaaccagg atttatacaa

1141 ggaggagaaa atgaaagcca tacgggaagc aatagcatga tacaaaggca ttaaagcagc

1201 gtatccacat agcgtaaaag gagcaacata gttaagaata ccagtcaatc tttcacaaat

1261 tttgtaatcc agaggttgat tatcgataag cttgatatcg aattcttact tgtacagctc

1321 gtccatgccg ccggtggagt ggcggccctc ggcgcgttcg tactgttcca cgatggtgta

1381 gtcctcgttg tgggaggtga tgtccaactt gatgttgacg ttgtaggcgc cgggcagctg

1441 cacgggcttc ttggccttgt aggtggtctt gacctcagcg tcgtagtggc cgccgtcctt

1501 cagcttcagc ctctgcttga tctcgccctt cagggcgccg tcctcggggt acatccgctc

1561 ggaggaggcc tcccagccca tggtcttctt ctgcattacg gggccgtcgg aggggaagtt

1621 ggtgccgcgc agcttcacct tgtagatgaa ctcgccgtcc tgcagggagg agtcctgggt

1681 cacggtcacc acgccgccgt cccctcagtt cattacggct ccaaggccta cgtgaagcac

1741 cccgccgaca tccccgacta cttgaagctg tccttccccg agggcttcaa gtgggagcgc

1801 gtgatgaact tcgaggacgg cggcgtggtg accgtgaccc aggactcctc cctgcaggac

1861 ggcgagttca tctacaaggt gaagctgcgc ggcaccaact tcccctccga cggccccgta

1921 atgcagaaga agaccatggg ctgggaggcc tcctccgagc ggatgtaccc cgaggacggc

1981 gccctgaagg gcgagatcaa gcagaggctg aagctgaagg acggcggcca ctacgacgct

2041 gaggtcaaga ccacctacaa ggccaagaag cccgtgcagc tgcccggcgc ctacaacgtc

2101 aacatcaagt tggacatcac ctcccacaac gaggactaca ccatcgtgga acagtacgaa

2161 cgcgccgagg gccgccactc caccggcggc atggacgagc tgtacaagta agaattcgat

2221 atcaagctta tcgataatca acctctggat tacaaaattt gtgaaagatt gactggtatt

2281 cttaactatg ttgctccttt tacgctatgt ggatacgctg ctttaatgcc tttgtatcat

2341 gctattgctt cccgtatggc tttcattttc tcctccttgt ataaatcctg gttgctgtct

2401 ctttatgagg agttgtggcc cgttgtcagg caacgtggcg tggtgtgcac tgtgtttgct

2461 gacgcaaccc ccactggttg gggcattgcc accacctgtc agctcctttc cggaactttc

2521 gctttccccc tccctattgc cacggcggaa ctcatcgccg cctgccttgc ccgctgctgg

2581 acaggggctc ggctgttggg cactgacaat tccgtggtgt tgtcggggaa atcatcgtcc

2641 tttccttggc tgctcgccta tgttgccacc tggattctgc gcgggacgtc cttctgctac

2701 gtcccttcgg ccctcaatcc agcggacctt ccttcccgcg gcctgctgcc ggctctgcgg

2761 cctcttccgc gtcttcgcct tcgccctcag acgagtcgga tctccctttg ggccgcctcc

2821 ccgcatcgat accgagcgct gctcgagaga tctacgggtg gcatccctgt gacccctccc

2881 cagtgcctct cctggccctg gaagttgcca ctccagtgcc caccagcctt gtcctaataa

2941 aattaagttg catcattttg tctgactagg tgtccttcta taatattatg gggtggaggg

3001 gggtggtatg gagcaagggg caagttggga agacaacctg tagggcctgc ggggtctatt

3061 gggaaccaag ctggagtgca gtggcacaat cttggctcac tgcaatctcc gcctcctggg

3121 ttcaagcgat tctcctgcct cagcctcccg agttgttggg attccaggca tgcatgacca

3181 ggctcagcta atttttgttt ttttggtaga gacggggttt caccatattg gccaggctgg

3241 tctccaactc ctaatctcag gtgatctacc caccttggcc tcccaaattg ctgggattac

3301 aggcgtgaac cactgctccc ttccctgtcc ttctgatttt gtaggtaacc acgtgcggac

3361 cgagcggccg caggaacccc tagtgatgga gttggccact ccctctctgc gcgctcgctc

3421 gctcactgag gccgcgcggg caaaggcccc ggcgccgggc gacctttgct cgcgcggcct

3481 cagtgagcga gcgagcgcgc agagagggag tggccaaa

//
